# Differential Gene Expression and Functional Enrichment Analysis of Breast Tumor and Normal Tissue RNA-seq Data: A Reanalysis of GEO Dataset GSE210787

**DOI:** 10.64898/2026.10.29.762989

**Authors:** Niteesh Surisetty, Bhupender Singh

## Abstract

**Background:** RNA sequencing (RNA-seq) allows a global assessment of expression differences between biological states. The present study reanalyzed publicly available RNA-seq data from human breast tumors and normal tissues to identify DEGs and to investigate functional enrichment.

**Methods:** Twelve paired-end RNA-seq samples from the GEO series GSE210787 were analyzed, including six tumor and six normal samples. Reads were obtained using the SRA Toolkit, checked with FastQC and MultiQC, trimmed with fastp, aligned to the GRCh38.p14 reference with STAR, and quantified at the gene level with featureCounts. Differential expression analysis was performed using DESeq2 in the Galaxy environment. The project report details criteria of adjusted p-value < 0.05 and absolute log2 fold change ≥ 1. Gene Ontology enrichment was performed using g:Profiler and KEGG overrepresentation analysis was performed separately for the up- and down-regulated gene sets.

**Results:** The project report contains a list of 120 differentially expressed genes (DEGs), including 69 upregulated and 51 downregulated genes in tumor tissue compared with normal tissue. One upregulated-gene GO Cellular Component term, cell periphery (GO:0071944), was observed with a P-value of approximately 0.017585 and 29 overlapping genes. The original report does not specify if this P-value is nominal or adjusted. No significant GO terms were found for the down-regulated gene set. After multiple testing corrections, no KEGG pathway was significant; the lowest adjusted p-value for the upregulated group was about 0.71393.

**Conclusion:** The project identified differences in expression between tumor and normal samples, but no KEGG pathway reached the set FDR significance threshold. The final interpretation needs to check sample metadata, DESeq2 contrast, GO significance statistic, KEGG background universe and analytical parameters against the original results.

## 1. Introduction

Molecular and cellular variations between tumors and between tumor and normal breast tissues define breast cancer as a heterogeneous disease. These variations may have an impact on the growth, development, and biological behavior of tumors. Finding the genes and biological mechanisms that differentiate tumor tissue from healthy tissue thus requires an understanding of the molecular alterations linked to breast cancer. High-throughput transcriptomic technologies allow the simultaneous analysis of thousands of transcripts and offer a more comprehensive picture of changes in gene expression, while traditional molecular biology methods can study individual genes.

A high-throughput sequencing technique called RNA sequencing (RNA-seq) is used to assess transcript abundance and look into variations in gene expression across biological conditions. RNA-seq can identify previously unidentified transcriptional changes and detect a wide range of expressed genes when compared to methods that look at a predetermined set of transcripts. Genes whose expression varies between experimental groups can be found using differential gene-expression analysis of RNA-seq data. However, proper sequencing quality control, read preprocessing, transcript quantification or reference-genome alignment, statistical analysis, and multiple hypothesis testing correction are necessary for accurate interpretation (Conesa et al., 2016). In cancer transcriptomics, where biological heterogeneity and technical variation can affect observed expression patterns, these steps are especially crucial..

Although a gene list by itself cannot completely explain the biological processes represented by the molecular alterations linked to breast tumors, differentially expressed genes can offer insights into those changes. By determining whether particular genes are overrepresented in recognized biological categories, functional enrichment analysis overcomes this restriction. While the Kyoto Encyclopedia of Genes and Genomes (KEGG) offers details on molecular pathways and biological systems, Gene Ontology (GO) analysis categorizes genes based on biological processes, molecular functions, and cellular components (Ashburner et al., 2000; Kanehisa & Goto, 2000). These methods can assist in classifying gene-expression data into functional themes that are easy to understand. However, because enrichment results rely on the input gene list, gene identifiers, background gene universe, and statistical correction for multiple comparisons, they should be interpreted with caution.

The current study made use of publicly accessible RNA-seq data from the Gene Expression Omnibus series GSE210787, which is linked to studies examining tumor-specific circular RNAs and their possible function in antitumor immune responses by generating cryptic peptides (Huang et al., 2024). Gene-level expression variations between breast tumor and normal tissue samples are the main focus of the current analysis. It does not directly test whether circular RNAs generate cryptic peptides or trigger immune responses, which sets it apart from the original study’s examination of circular RNA biology. Rather, it makes use of the available sequencing data to investigate the functional properties of the resulting gene sets as well as variations in gene expression between the two tissue groups.

In this study, quality assessment, read preprocessing, reference-genome alignment, and gene-level read counting were applied to paired-end RNA-seq data from six tumor and six normal samples. Next, genes with changed expression between tumor and normal samples were found using differential expression analysis. To look into possible functional relationships between upregulated and downregulated gene sets, separate analyzes of KEGG pathway overrepresentation and Gene Ontology enrichment were performed. Characterizing tumor-versus-normal gene-expression differences, summarizing the functional categories linked to the chosen genes, and determining whether any KEGG pathways remained significant after multiple-testing correction were the goals. This study offers a targeted exploratory evaluation of transcriptomic variations in the examined breast tissue samples by combining differential expression analysis with functional enrichment.

## 2. Materials and Methods

### 2.1. Dataset and sample design

The source dataset is identified in the project report as GEO series GSE210787. Twelve Homo sapiens samples—six tumor and six normal samples—that were produced using an Illumina NovaSeq 6000 platform made up the recorded design. The report’s sample-to-run mapping is displayed below. Prior to submission, this mapping needs to be verified against the GEO/SRA metadata.

#### Sample and SRA run organization

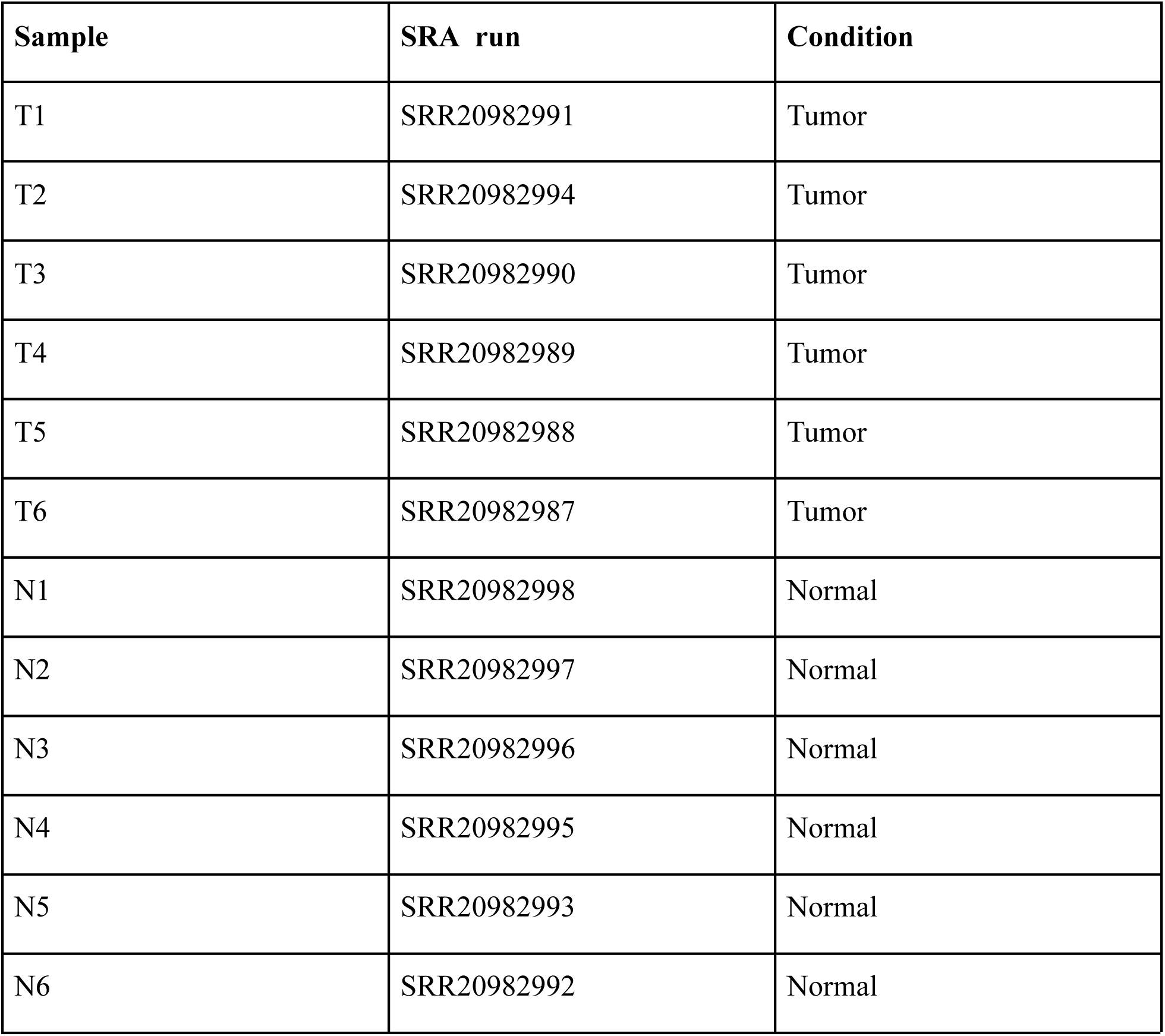

### 2.2. Reference genome and annotation

The use of the human GRCh38.p14/hg38 reference genome and GENCODE Release 50 annotation is documented in the project report. FASTA and GENCODE v50 GTF were utilized as the primary assemblies. According to the report, compatibility of chromosome names was verified prior to alignment and counting.

### 2.3. Read retrieval and quality control

SRA Toolkit 3.2.1(NCBI Official documentation) was used to retrieve the 12 SRA runs as paired-end FASTQ(Andrews,2020), data. MultiQC 1.21(Ewels et al., 2016) was utilized to combine quality-control summaries, and FastQC 0.12.1 was utilized for raw-read and post-trimming quality evaluation. According to the project report, GC content in inspected samples ranges from 49 to 52%, per-sequence quality is generally acceptable, overrepresented sequences are less than 1% in reported samples, and adapter content is acceptable after trimming. Per-base sequence-content warnings and sequence duplication of roughly 50–60% were found in numerous samples. Rather than treating these metrics as independent proof of failure, the report correctly views them as observations that need to be interpreted in the context of RNA-seq libraries.

### 2.4. Read trimming

Fastp version 1.0.1(Chen et al., 2018) was used to process paired-end reads in order to eliminate sequencing adapters and low-quality read segments. All 12 samples were successfully preprocessed, and both read mates were kept, according to the report. Quality control was then carried out after trimming.

### 2.5. Alignment and gene-level quantification

When the project first tried to generate STAR(Dobin et al., 2013) genome-indexes in WSL/Ubuntu, it kept stalling during junction insertion due to the limited local resources. The alignment stage was shifted to Galaxy on the advice of the mentor. According to the report, 12 BAM outputs were produced using STAR version 2.7.11b+galaxy1 with the hg38 reference and GENCODE v50 annotation. Using exon features and the gene_id attribute, gene-level quantification was carried out using featureCounts version 2.1.1+galaxy1(Liao et al., 2014). Reads with paired ends were tallied as fragments. The original Galaxy history should be used to confirm the precise featureCounts strandedness and other parameters.

### 2.6. Differential expression analysis

The Galaxy DESeq2 tool (version 2.11.40.8+galaxy4) was used to perform differential expression. Tumor versus normal was defined as the condition factor. The entire DESeq2(Love et al., 2014), result was kept as Galaxy dataset #145. The project report documents the selection criteria of absolute log2 fold change ≥ 1 and adjusted p-value (padj) < 0.05. Before publishing, the saved Galaxy(Afgan et al.,2018) run must be examined for the precise design formula, reference level, contrast direction, and filtering settings.

### 2.7. Visualization and functional enrichment

The project report details a heatmap based on variance-stabilizing-transformation (VST) expression values for the 120 chosen DEGs and a volcano plot created from the full DESeq2 result. Additionally, a PCA plot was produced. Using g:Profiler GOSt version 0.1.7+galaxy11, Gene Ontology enrichment was carried out independently for upregulated and downregulated genes across Biological Process, Molecular Function, and Cellular Component. To increase identifier compatibility, Ensembl version suffixes were eliminated. After converting Ensembl identifiers to Entrez identifiers and eliminating unmapped entries, KEGG overrepresentation analysis was carried out independently for upregulated and downregulated gene sets using a specialized Galaxy KEGG ORA tool.

## 3. Results

### 3.1. Quality control and preprocessing

Twelve biological samples, represented by twenty-four paired-end FASTQ files, were processed, according to the report. The examined quality-control summaries revealed GC distributions primarily spanning roughly 45–65% in the MultiQC visual summary and per-sequence quality around Phred 33 in the displayed plot, whereas the narrative reports sample GC content around 49–52%. The MultiQC heatmap that was displayed was primarily green for adapter and overrepresented-sequence statuses. Samples differed in terms of sequence duplication, and precise percentages were not verified. For certain samples, per-base sequence-content warnings were visible.

### 3.2. Alignment and gene-level quantification

According to the report, 12 BAM outputs were generated and the local STAR indexing limitation was overcome by shifting the alignment step to Galaxy. For each of the 12 samples, featureCounts produced gene-level count tables that were fed into DESeq2. Alignment rates and feature-assignment percentages should not be added unless verified by the original output files because they are not expressed numerically in the report text.

### 3.3. Differentially expressed genes

The project report finds 120 DEGs, 69 of which are upregulated and 51 of which are downregulated in the tumor group, using the recorded criteria of adjusted p-value < 0.05 and |log2 fold change| ≥ 1. The PCA and heatmap, according to the report, demonstrate different expression patterns between the tumor and normal groups.

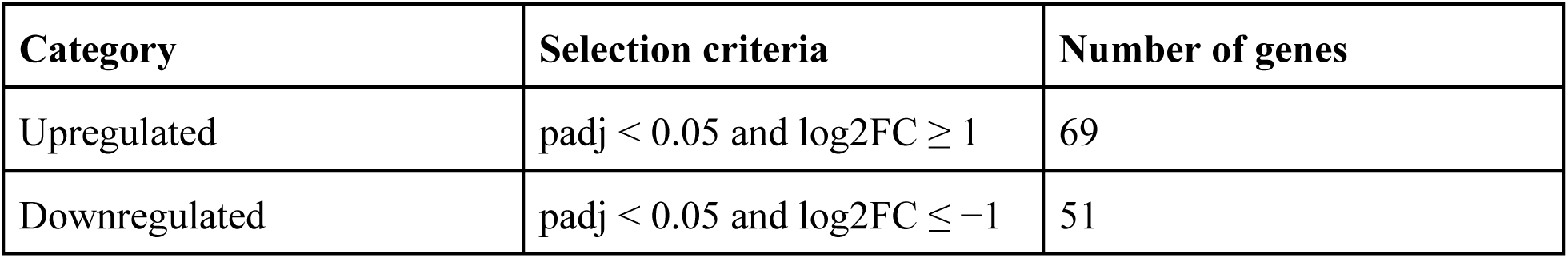
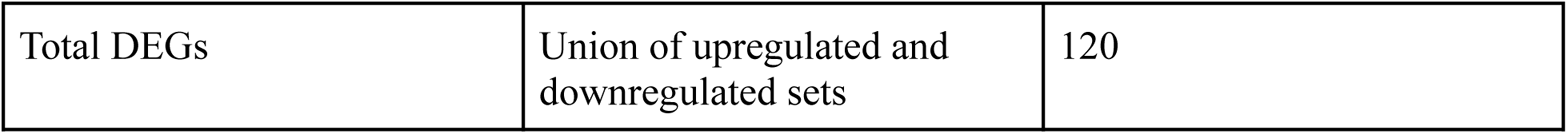

| **Category** | **Selection criteria** | **Number of genes** |
|---|---|---|
| Upregulated | padj < 0.05 and log2FC ≥ 1 | 69 |
| Downregulated | padj < 0.05 and log2FC ≤ −1 | 51 |
| Total DEGs | Union of upregulated and downregulated sets | 120 |

**Figure 1.**
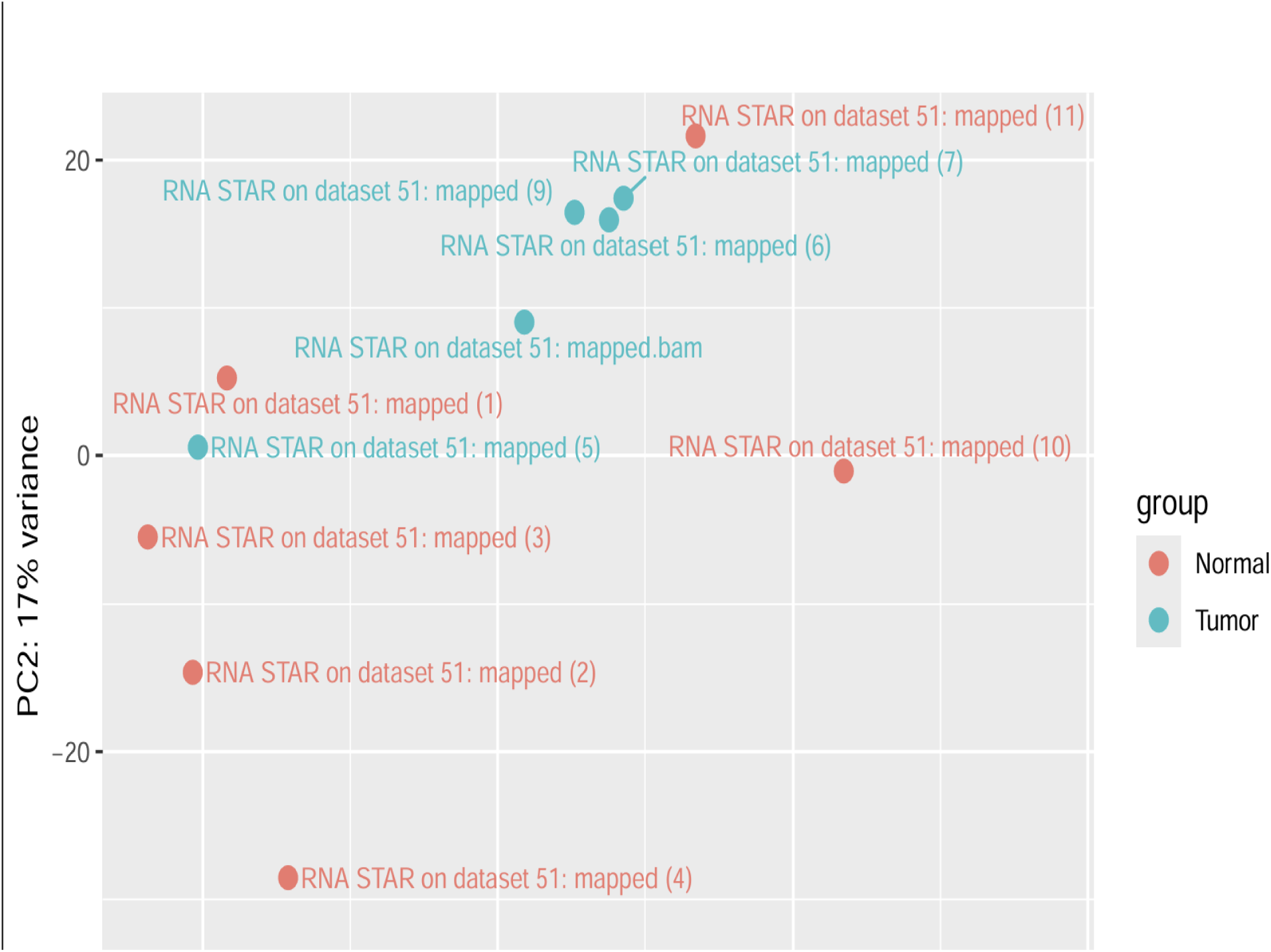
Principal component analysis (PCA) of breast tumor and normal samples.

PCA was used to compare and contrast the global gene-expression profiles of tumor and normal samples using variance-stabilized gene-expression data. One sample is represented by each point. The legend indicates that blue-green points indicate tumor samples and red points indicate normal samples. 17% of the variance is explained by PC2. After verifying its value from the initial plot or analysis output, the percentage of variance explained by PC1 should be included.

**Figure 2.**
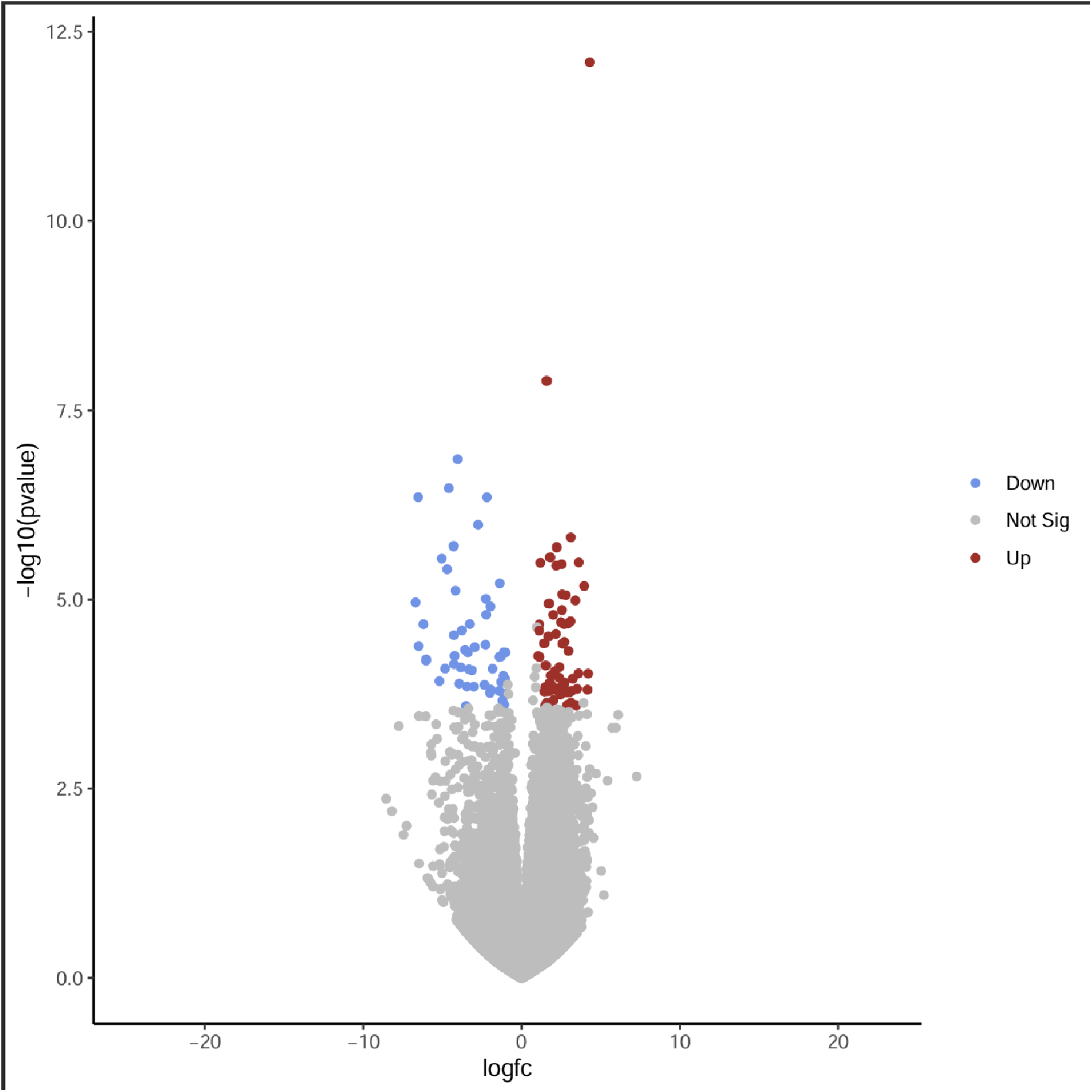
Volcano plot showing differential gene expression between breast tumor and normal samples.

The y-axis shows −log₁₀ of the nominal p-value, and the x-axis shows log₂ fold change (log₂FC). A gene examined in the differential expression analysis is represented by each point. Gray points show genes that did not fit the criteria for differential expression, red points show upregulated genes, and blue points show downregulated genes. An absolute log2 fold change of at least one and an adjusted p-value (padj) of less than 0.05 were used to classify genes. The categories of differential expression are indicated by the color legend.

**Figure 3.**
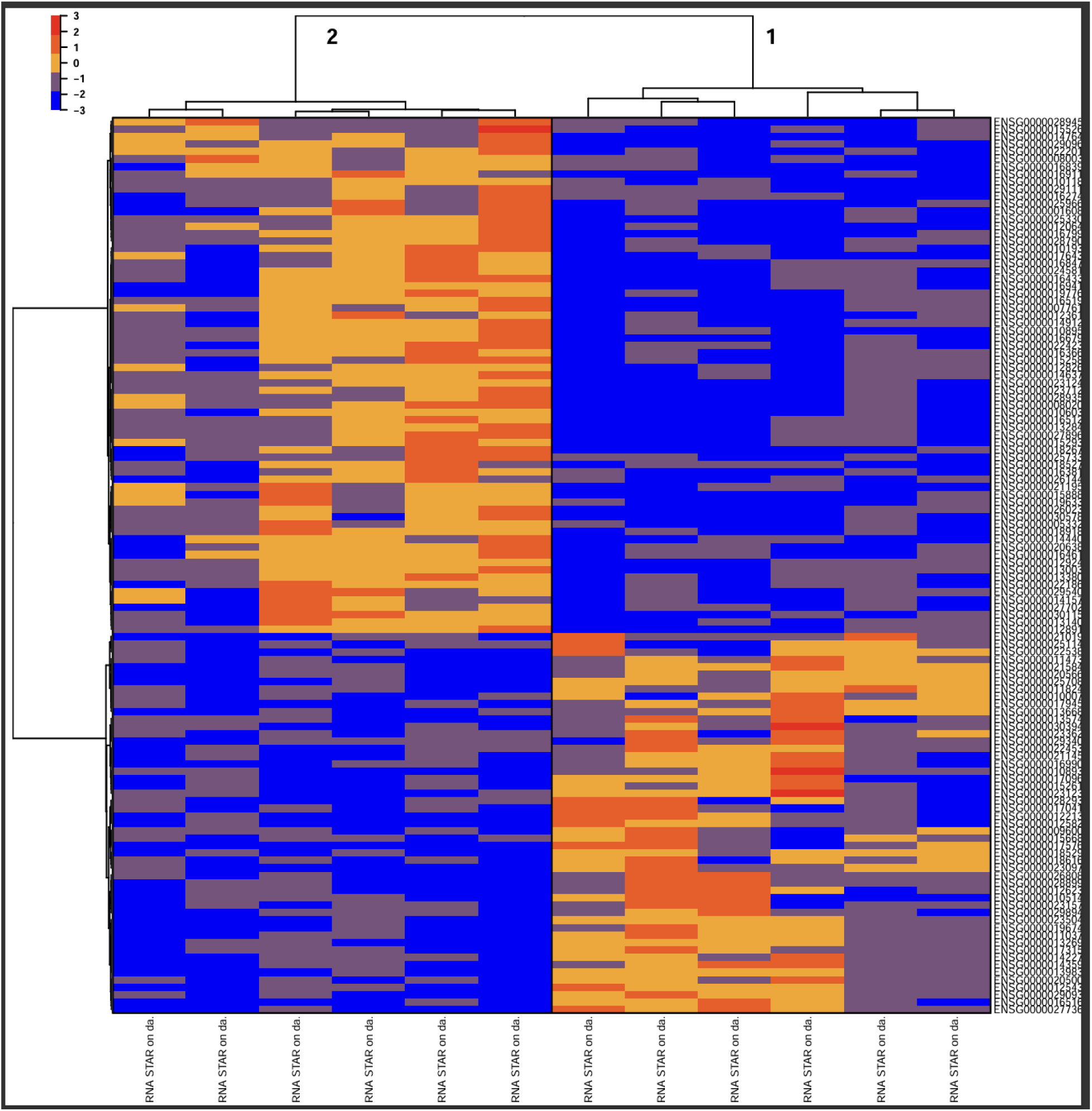
Hierarchical clustering heatmap of gene expression across tumor and normal samples.

The heatmap shows the relative levels of gene expression in each of the 12 samples (6 tumor and 6 normal). A gene is represented by each row, and a sample is represented by each column. According to the displayed scale, the color scale represents relative expression, with blue denoting lower expression and yellow to red denoting higher expression. Genes and samples are grouped using hierarchical clustering based on similarities in their expression patterns.

### 3.4. Gene Ontology enrichment

The report lists 29 genes in the intersection and one GO Cellular Component term, cell periphery (GO:0071944), with a p-value of roughly 0.017585 for the upregulated gene set. For the downregulated gene set, no significant GO term was found. Verify whether 0.017585 is the nominal p-value or the multiple-testing-adjusted value from the original g:Profiler(Raudvere et al., 2019) result prior to publication, and clearly report the relevant statistic.

| **Gene set** | **Reported GO finding** | **Evidence recorded** |
|---|---|---|
| Upregulated | GO:0071944—cell periphery (Cellular Component) | p ≈ 0.017585; 29 intersecting genes; confirm nominal vs adjusted p-value |
| Downregulated | No significant GO term reported | No term passed the selected significance setting, as recorded in the report |

### 3.5. KEGG pathway enrichment

Following the conversion of Ensembl-to-Entrez identifiers and the elimination of unmapped identifiers, KEGG(Kanehisa & Goto, 2000), overrepresentation analysis was carried out independently for upregulated and downregulated gene lists. According to the report, after multiple-testing correction, none of the pathways that had nominal p-values below 0.05 were still significant. Adjusted p-values for the downregulated set were reported as 1.0, while the lowest recorded adjusted p-value for the upregulated set was roughly 0.71393. Therefore, the reported adjusted p-value threshold of 0.05 was not met by any KEGG pathway.

| **Gene set** | **FDR-significant pathways** | **Recorded result** |
|---|---|---|
| Upregulated | 0 | Minimum reported adjusted p-value ≈ 0.71393 |
| Downregulated | 0 | Top adjusted p-values reported as 1.0 |

The reported minimum adjusted p-value of roughly 0.71393 would not be significant if the significance cutoff were only changed to 0.10 or 0.20. If the input gene list or other analysis parameters were altered for a reason supported by science, a new enrichment result might be different; adjusted p-values should not be manually modified. Any sensitivity analysis ought to be repeated using the initial findings and recorded.

**Table 4.** KEEG pathway over-representation analysis of upregulated genes.

| **KEGG pathway** | **KEGG ID** | **Overlapping genes** | **Pathway size** | **Adjusted p-value** |
|---|---|---|---|---|
| Carbohydrate digestion and absorption | hsa04973 | 2 | 48 | 0.71393 |
| Integrin signaling | has04518 | 3 | 154 | 0.71393 |
| IgSF CAM signaling | hsa04517 | 4 | 300 | 0.71393 |
| Endocrine and other factor-regulated calcium reabsorption | hsa04961 | 2 | 53 | 0.71393 |
| Cell adhesion molecule (CAM) interaction | hsa04514 | 3 | 160 | 0.71393 |
| cGMP-PKG signaling pathway | hsa04022 | 3 | 166 | 0.71393 |
| Focal adhesion | hsa04510 | 3 | 203 | 0.99521 |
| ECM-receptor interaction | hsa04512 | 2 | 89 | 0.99521 |
| Complement and coagulation cascades | hsa04610 | 2 | 89 | 0.99521 |

Several pathways with nominal p-values less than 0.05 were found by KEGG pathway overrepresentation analysis of the upregulated gene set, including integrin signaling, IgSF CAM signaling, and carbohydrate digestion and absorption. After multiple-testing correction, none of them were still statistically significant (adjusted p-value greater than 0.05).

**Table 5.**
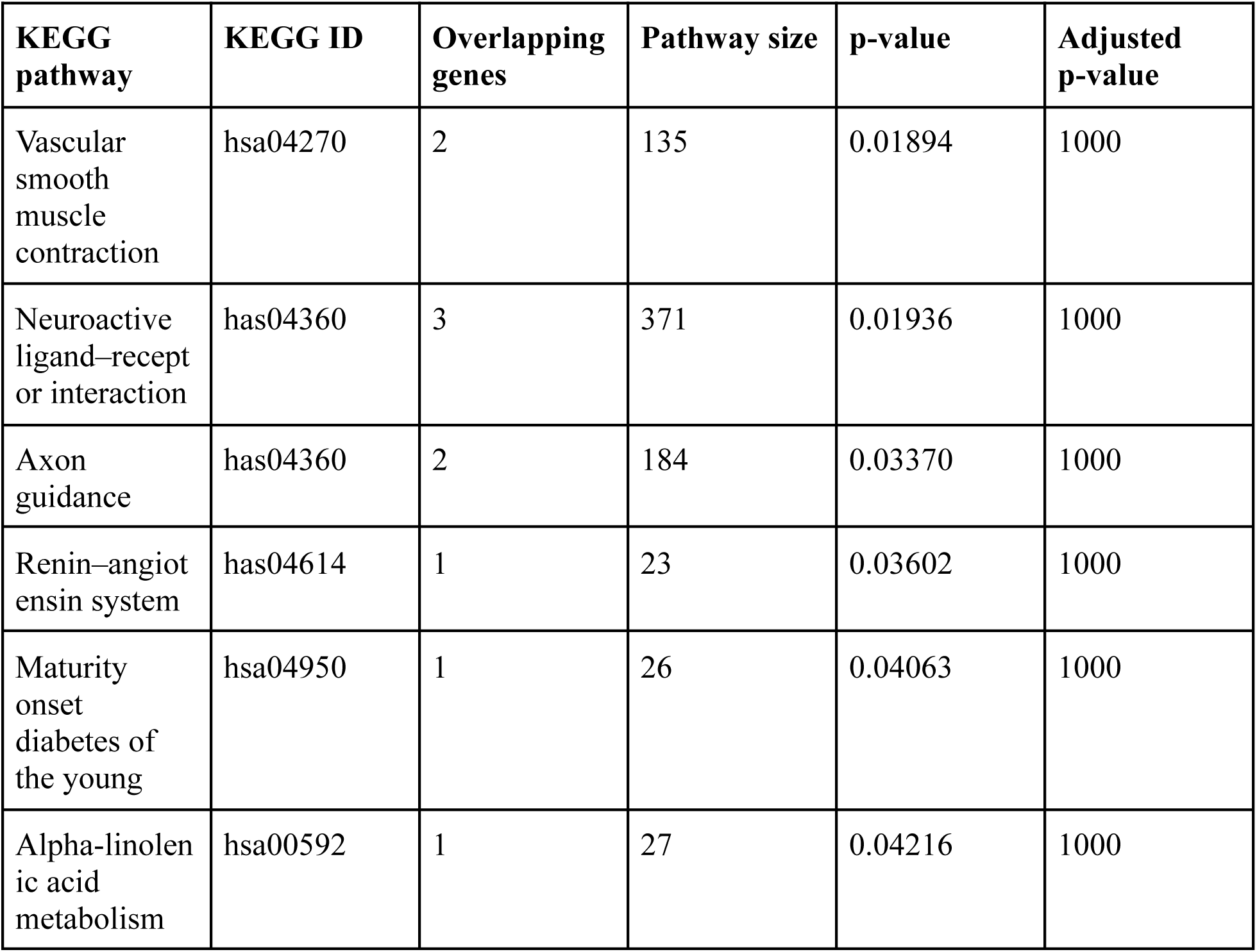
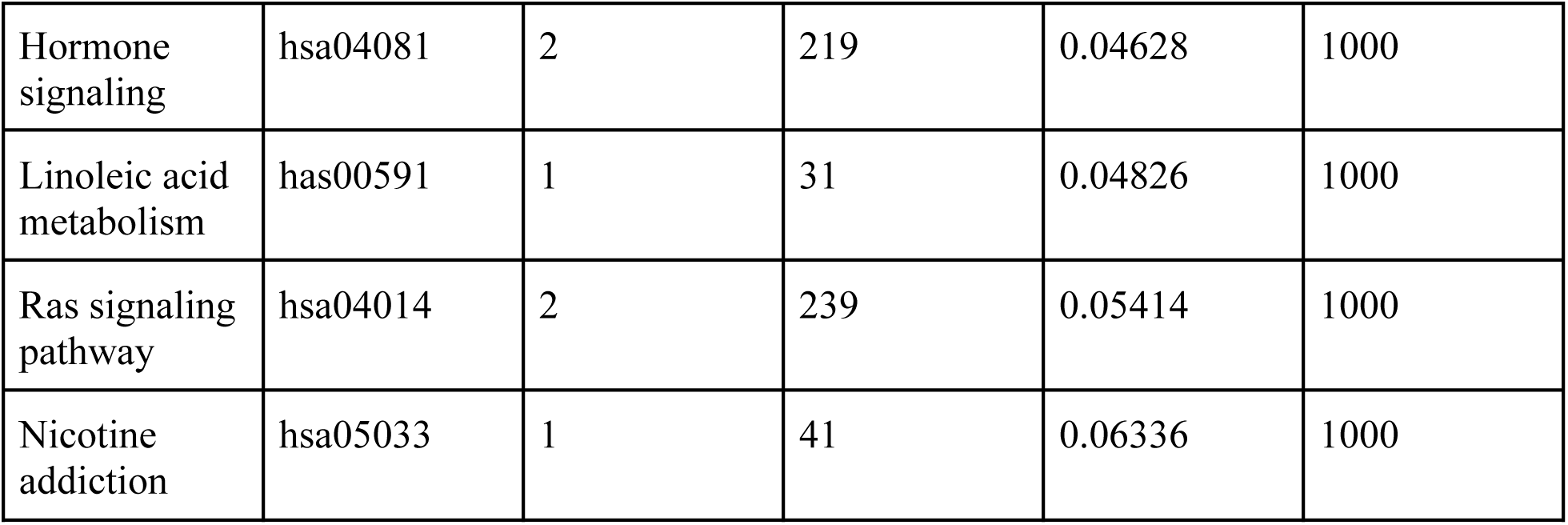
KEGG pathway over-representation analysis of downregulated genes.

| **KEGG pathway** | **KEGG ID** | **Overlapping genes** | **Pathway size** | **p-value** | **Adjusted p-value** |
|---|---|---|---|---|---|
| Vascular smooth muscle contraction | hsa04270 | 2 | 135 | 0.01894 | 1000 |
| Neuroactive ligand–receptor interaction | has04360 | 3 | 371 | 0.01936 | 1000 |
| Axon guidance | has04360 | 2 | 184 | 0.03370 | 1000 |
| Renin–angiotensin system | has04614 | 1 | 23 | 0.03602 | 1000 |
| Maturity onset diabetes of the young | hsa04950 | 1 | 26 | 0.04063 | 1000 |
| Alpha-linolenic acid metabolism | hsa00592 | 1 | 27 | 0.04216 | 1000 |
| Hormone signaling | hsa04081 | 2 | 219 | 0.04628 | 1000 |
| Linoleic acid metabolism | has00591 | 1 | 31 | 0.04826 | 1000 |
| Ras signaling pathway | hsa04014 | 2 | 239 | 0.05414 | 1000 |
| Nicotine addiction | hsa05033 | 1 | 41 | 0.06336 | 1000 |

The downregulated gene set was subjected to KEGG pathway overrepresentation analysis. Vascular smooth muscle contraction (p = 0.01894), neuroactive ligand–receptor interaction (p = 0.01936), and axon guidance (p = 0.03370) were among the pathways with the lowest nominal p-values. Nevertheless, all reported adjusted p-values were 1.000, meaning that none of the pathways were statistically significant after multiple-testing correction. As a result, rather than being proof of substantial pathway enrichment, these findings should be viewed as exploratory observations.

## 4. Discussion

In this study, we used the GSE210787 dataset to analyze RNA-seq data from breast tumor and normal samples. Finding genes that expressed differently in the two groups and learning about their potential biological roles was the primary goal. 120 differentially expressed genes (DEGs), including 69 upregulated and 51 downregulated genes in tumor samples, were reported based on the analysis’s criteria. These findings offer a preliminary understanding of the variations in gene expression between the two groups. However, since only six tumor samples and six normal samples were included, the results should be interpreted cautiously. The reliability of RNA-seq results is influenced by the study design, sample quality, data processing, and statistical techniques, as explained by Conesa et al. (2016).

The sample’s overall expression patterns were examined using the PCA plot and heatmap. Plots of the tumor and normal groups were displayed. The variations revealed by the differential expression analysis were in line with these patterns. Plots by themselves, however, are unable to verify the biological function of individual genes or explain why the samples differ. Because DESeq2 takes sample variation into account when estimating changes in gene expression, it is frequently used to analyze RNA-seq count data (Love et al., 2014). The reported DEG list in this study was predicated on an absolute log2 fold change of at least 1 and an adjusted p-value less than 0.05. Before publication, the final number of genes should be verified using the original DESeq2 output.

I used Gene Ontology (GO) enrichment analysis to comprehend the potential roles of the chosen genes. The cell periphery term (GO:0071944) in the upregulated gene set was found to have 29 intersecting genes and a p-value of roughly 0.017585. Cellular elements at or close to the cell boundary are included in this term. Because the result identifies genes related to the cell periphery, it might be helpful for future research. Before determining whether this term is statistically significant, I must ascertain whether the reported p-value is the original p-value or the adjusted p-value. For the downregulated genes, no noteworthy GO terms were found. GO analysis aids in classifying genes based on recognized biological functions; however, the outcomes rely on the genes that are submitted, the annotations that are available, and the statistical method used (Ashburner et al., 2000; Raudvere et al., 2019).

Additionally, I conducted separate KEGG pathway analyzes for the genes that were upregulated and downregulated. After multiple-testing correction, none of the pathways that had nominal p-values below 0.05 remained significant. The downregulated gene set’s adjusted p-value was 1.0, while the upregulated gene set’s lowest adjusted p-value was roughly 0.71393. As a result, there is insufficient statistical evidence in the current results to classify any KEGG pathway as significantly enriched. The results for upregulated genes included pathways associated with integrin signaling and cell adhesion, but they cannot be considered significant findings in this analysis. Although KEGG is a helpful tool for researching established biological pathways, the pathway results must be interpreted using both the biological data and the corrected p-values (Kanehisa) & Goto, 2000).

The comparatively small number of genes in the chosen lists could be one explanation for not discovering significant KEGG pathways. The selection of background genes and the conversion of gene identifiers may also have an impact on the enrichment results. Furthermore, multiple-testing correction may make it more difficult for a pathway to maintain statistical significance because the analysis tests numerous pathways. When analyzing and interpreting RNA-seq data, these are typical considerations (Conesa et al., 2016). Because of this, the pathways with lower nominal p-values should not be presented as verified biological findings, but they can be thought of as potential areas for additional research. Additionally, the KEGG analysis’s gene identifiers and background gene set should be compared to the original analysis settings.

A published study on tumor-specific circular RNAs and their function in anti-tumor immunity is linked to the dataset utilized in this work (Huang et al., 2024). Circular RNAs capable of producing cryptic peptides and their potential impact on immune responses were examined in that study. The focus of my analysis was different: I used functional enrichment analysis to compare the gene-level expression of the tumor and normal samples. Circular RNA expression, peptide synthesis, and the immune mechanisms discussed in the original paper were not directly examined by me. As a result, my findings should not be interpreted as direct confirmation of the mechanisms described by Huang et al. (2024), but rather as a distinct analysis of the dataset.

This study has certain limitations. First, a larger dataset would be helpful to determine whether the same expression patterns are seen in additional samples, as the analysis only included 12 samples. Second, the settings used for differential expression analysis and the accurate assignment of samples to the tumor and normal groups determine the outcomes. Third, no KEGG pathway met the reported adjusted p-value threshold, and the GO p-value must be verified for statistical significance. Lastly, I did not validate the identified genes through independent experiments. When interpreting the results, these factors should be taken into account. Drawing trustworthy conclusions from RNA-seq analyzes requires careful study design and validation, as explained by Conesa et al. (2016).

Next, I would check the GO and KEGG output tables, the sample data, and the DESeq2 results. After that, I would think about examining the significant genes in a separate dataset to see if there are any comparable changes in expression. An alternative would be to carry out a ranked-gene analysis, like gene set enrichment analysis, which looks for trends throughout a ranked list of genes instead of relying solely on a chosen DEG list. Investigating whether gene groups exhibit coordinated changes even when individual pathways fall short of the current significance threshold could be aided by this. But since this would be a fresh analysis, its findings would have to be evaluated independently.

overall, this study found documented variations in gene expression between breast tumor and normal samples and offered a preliminary functional analysis of the chosen genes. Although its significance needs to be verified, the GO result pertaining to the cell periphery might be worth investigating further. After multiple-testing correction, no statistically significant pathway was found in the KEGG analysis. As a result, it is best to view the current results as a first step toward comprehending the variations in gene expression in these samples; more research and validation are required before making more robust biological inferences.

## Supporting information

https://docs.google.com/document/d/1rp53leZwKXKNVdDA8XTg7dEvzen0BYbzMTa0Xz1lD48/edit?tab=t.0

## Data availability

The GEO series GSE210787 and associated SRA records make the source sequencing dataset accessible to the general public. The final count matrix, full DESeq2 results, enrichment tables, analysis settings, and any reproducible scripts or workflow exports that are genuinely accessible and authorized for sharing should be provided or deposited prior to manuscript submission. Don’t announce the availability of additional files until they’ve been put together and examined.

## Ethics, permissions, and authorship

Secondary analysis of publicly accessible RNA-sequencing data from the Gene Expression Omnibus (GEO) under accession GSE210787 was used in this study. No additional biological samples were gathered for this analysis, nor were any new human subjects recruited. Publicly accessible sequencing data was used in the investigation. The policies of the target journal and the relevant institution should be followed when determining whether further ethical approval is required.

## Notes

https://www.ncbi.nlm.nih.gov/geo/query/acc.cgi?acc=GSE210787

### Competing Interest Statement

The authors have declared no competing interest.

