## Supplementary material for "Differential Gene Expression and Functional Enrichment Analysis of Breast Tumor and Normal Tissue RNA-seq Data: A Reanalysis of GEO Dataset GSE210787": https://docs.google.com/document/d/1rp53leZwKXKNVdDA8XTg7dEvzen0BYbzMTa0Xz1lD48/edit?tab=t.0

A Project Report on

### RNA-SEQ DIFFERENTIAL EXPRESSION ANALYSIS:IDENTIFICATION OF DIFFERENTIALLY EXPRESSED GENES BETWEEN BREAST TUMOR AND NORMAL TISSUE


**Submitted By:**

**Surisetty Niteesh**

Next-Gen Sequencing & Multi-Omics Data Analysis

2 Months

**PI Details:**

**DR.Bhupender Singh**

CRO Scientist

Biotecnika Info Lab Pvt.Ltd


**OCTOBER 2026**

**DECLARATION**

I, ‘Surisetty Niteesh’, hereby declare that the project work entitled “RNA-Seq Differential Expression Analysis” submitted to Biotecnika Info Labs is an original work carried out by me under the guidance of Dr. Bhupender Singh.

I declare that this project has not been submitted to any other University or Institution for the award of any degree or diploma or any other title.

Further, I declare that I shall not publish, present, or communicate any part of this project work in any form– including journals, conferences, or digital media, without the prior written consent of my Principal Investigator and the host organization.

I understand that failure to obtain such prior approval shall be treated as a breach of institutional and research ethics, and may result in disciplinary action as deemed appropriate by the organization.
By signing below, I acknowledge that I have read and agree to abide by the above statement.

.

**Student Signature:** 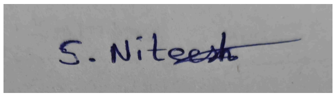

**Student Name:** Surisetty Niteesh
**Course Name:** Next-Gen Sequencing & Multi-Omics Data Analysis
**Project Duration:** 2 Months

Dr Bhupender Singh
Scientist & Trainer
Biotecnika Info Labs Pvt Ltd

### ACKNOWLEDGEMENT

I would like to express my sincere gratitude to Dr. Bhupender Singh, Scientist & Trainer at Biotecnika Info Labs Pvt. Ltd., for his guidance throughout this project. His emphasis on following a structured RNA-seq workflow, documenting each step, and checking results before moving to the next stage helped me understand that bioinformatics analysis is not simply a sequence of software commands, but a process in which biological reasoning and computational quality control have to work together.

I am grateful for the opportunity to work with a real public RNA-seq dataset and to gain practical experience with quality control, read preprocessing, genome alignment, gene-level quantification, differential expression analysis, visualization, and functional enrichment. The project also gave me experience in troubleshooting computational problems, particularly when the local STAR genome-indexing step became impractical and the analysis had to be moved to Galaxy without changing the intended analytical strategy.

Finally, I thank everyone who supported me during the training period and contributed directly or indirectly to the completion of this project.

#### TABLE OF CONTENTS

##### List of figures


Figure 1. Dataset/GEO. [Page.no 12]
Figure 2. Reference Genome and Annotation Download. [Page.no 15]
Figure 3. RNA-Seq Dataset and Trimmed Reads. [Page.no 16]
Figure 4. FastQC Quality Control Report. [Page.no 17]
Figure 5. FastQC Analysis of Trimmed Reads. [Page.no 18]
Figure 6. STAR Alignment Configuration. [Page.no 19]
Figure 7. featureCounts Configuration. [Page.no 20]
Figure 8. DESeq2 Parameters. [Page.no 21]
Figure 9. Volcano Plot of Differentially Expressed Genes. [Page.no 22]
Figure 10. Heatmap Analysis Results. [Page.no 23]
Figure 11. GO Functional Enrichment Analysis. [Page.no 24]
Figure 12. Per-Base Sequence Content and GC Content. [Page.no 26]
Figure 13. STAR Alignment BAM Output. [Page.no 27]
Figure 14. featureCounts Gene Count Results. [Page.no 28]
Figure 15. DESeq2 Differential Expression Analysis. [Page.no 29]
Figure 16. DESeq2 PCA Plot. [Page.no 30]
Figure 17. GO Enrichment Analysis Results. [Page.no 32]
Figure 18,19 .KEGG Pathway Enrichment Results Both Upregulated & Downregulated. [Page.no 34]
Figure 20. KEGG Enrichment Filtering Results. [Page.no 42]


##### List of Tables


Table 1. Dataset and experimental design. [Page.no 11]
Table 2. Sample-to-SRA run mapping maintained throughout the analysis. [Page.no 12]
Table 3. Software, versions and purposes recorded. [Page.no 13]
Table 4. Differential-expression selection criteria. [Page.no 14]
Table 5. Summary of quality-control observations. [Page.no 24]
Table 6. Final differential-expression counts. [Page.no 28]
Table 7. GO enrichment summary. [Page.no 29]
Table 8. KEGG pathway enrichment summary. [Page.no 32]
Table 9. Problems encountered and their resolution. [Page.no 34]
Table 10. Final outputs retained for the project report. [Page.no 35]

**ABSTRACT**

The analysis workflow encompassed all stages starting with the acquisition of public data and organization of samples, quality assessment using (RNA-seq) provides an overview of transcriptional activity on an entire genome scale and is being extensively utilized to discover molecular distinctions among various biological conditions. The present project is an application of the complete process for analysis of bulk RNA-seq data employing the publicly available breast tumor and normal tissue data derived from GEO accession *GSE210787.* The studyRNA analyzed *12 human RNA-seq samples* isolated from both tumor and normal tissues using the *Illumina NovaSeq 6000 technology*, with an equal number of tumor and normal samples.
g FastQC and MultiQC techniques, adapter and quality filtering using fastp, reference generation via GRCh38.p14 and GENCODE Release 50, splice aware alignment using STAR software package, gene quantification using featureCounts and differential expression analysis with DESeq2 program. Due to the recurrent delays encountered while performing local STAR genome indexing during junction insertion under WSL, it was advisable to use Galaxy for alignment step according to the recommendations of the project mentor. The use of Galaxy provided a successful result in the form of aligned BAM files for 12 samples .

We identified 120 differentially expressed genes (DEGs) in tumor samples versus normal samples using a modified p-value < 0.05 and a minimum absolute log2 fold-change of 1. Of these, 69 genes were over-expressed and 51 genes were under-expressed in the tumor samples. Principal component analysis and a heatmap of VST-normalized expression data revealed that the two groups had different expression patterns. Then, functional enrichment analysis was performed based on Gene Ontology (GO) and KEGG pathway overrepresentation analysis.

The elevated gene collection produced one statistically significant GO Cellular Component term, *GO:0071944 (cell periphery)*, with an intersection of 29 genes and a reported p-value of about 0.017585. The downregulated gene set showed no appreciable GO enrichment under the selected parameters. KEGG analysis came after the Ensembl-to-Entrez identification conversion. Despite the observation of nominal relationships, no KEGG pathway remained significant after multiple-testing correction in either direction. Overall, the experiment shows how important statistical correction is, how non-significant pathway results should be presented clearly, and how RNA sequencing can be repeated from raw sequencing data to differential expression and functional interpretation.

### INTRODUCTION

#### Background

RNA is an important intermediary molecule between genome and cell. RNA profiling is a technique that measures the expression levels of thousands of RNAs, giving insight into how cells respond to illness, growth, treatment, environmental conditions, or any other biological state. RNA-seq has emerged as a popular transcriptomic technology owing to its ability to quantify diverse sets of transcripts and be employed for various purposes including gene expression, transcript quantification, alternative splicing, and more. The reviews of RNA-seq analysis highlight the fact that not only the experiment but also quality control, pre-processing, alignment or quantification, statistical analysis and data interpretation are important in deriving biological inferences (Conesa et al., 2016; Qi et al., 2017).

In cancer research, comparison of cancerous and normal tissues may potentially provide the identification of transcriptional differences associated with the development of cancer and changed cellular environment. But from a biological point of view it may be difficult to interpret a simple list of differentially regulated genes. Gene functional annotation and pathway analysis allow us to put information about genes within the framework of other biological systems. KEGG links genes and molecular activities to pathways, and Gene Ontology provides controlled vocabularies for biological processes, molecular functions and cellular components (Ashburner et al., 2000; Kanehisa & Goto, 2000).

#### Biological context of the selected dataset

In this project, we used the GEO series for analysis as *GSE210787, "Tumor-specific circRNAs elicit anti-tumor immune response via encoding cryptic peptides [RNA-Seq].* GEO description: experiment on expression profiling using *Homo sapiens* related to breast cancer and normal tissues, with *12 samples* prepared with Illumina NovaSeq 6000. The research article associated with this GEO series was focused on the tumor-specific circRNAs and their contribution to the generation of cryptic antigenic peptides and anti-tumor immune response. However, the present project is not aimed at replication of all the analyzes made in that study.

#### Rationale for the computational workflow

The RNA-seq analysis pipeline should not be considered as individual software tasks, but as a series of interconnected quality control steps. Prior to any kind of pre-processing, it is necessary to perform quality control on the raw reads, quality control on the trimmed reads, align or quantify the reads against the correct reference genome and analyze the count data using statistics specific for count data generated by sequencing. This pipeline is in line with that logic and current RNA-seq best practice guidelines (Conesa et al., 2016).

Thus, the pipeline ranged from data acquisition and quality control, to pre-processing, reference-based mapping, gene counts, differential expression, visualization and enrichment. . The pipeline was run partly in *WSL/Ubuntu and partly in Galaxy.* This approach enabled the computationally intensive STAR alignment step to be completed while keeping the mentor's preferred pipeline.

### REVIEW OF LITERATURE

#### 1. RNA-seq as a transcriptomic approach

The advent of RNA sequencing technology has revolutionized the study of transcriptomes by enabling simultaneous sequencing and quantitation of large populations of RNA molecules. The modern pipeline in RNA sequencing can be used for many different applications such as gene and transcript expression, alternative splicing, fusions and many more. The standard pipeline in studies on differential expression includes quality assessment, pre-processing, alignment or quantification of transcripts, summarization of counts, modeling and visualization (Conesa et al., 2016; Chowdhury et al., 2020).

#### 2. Quality control and preprocessing

The FastQC software's objective was to give a quick summary of the sequencing quality and any issues pertaining to high throughput sequencing data. The tool's module-based output assesses sequence content, GC content, duplication levels, overrepresented sequences, adapter content, and per base sequence quality. It might be beneficial to carry out the technical issues before proceeding with further analysis because some of them could affect how the findings of later analyzes are interpreted.

Fastp was selected to do adapter trimming and quality filtering in this specific instance. Fastp is an all-inclusive program that enables concurrent quality monitoring, adapter trimming, filtering, and multiple other activities (Chen et al., 2018). The data from several samples were summarized using MultiQC. MultiQC was specifically created to provide an overview of the outcomes generated by many tools and samples in a single report (Ewels et al., 2016).

#### 3. Reference alignment and read summarization

Specifically designed to swiftly map RNA-seq reads to a reference genome by accounting for spliced alignments, STAR is a splice-aware RNA-seq aligner. STAR is a suitable method since the aligned RNA-seq reads will be aggregated to the gene level in a traditional reference-based RNA-seq pipeline (Dobin et al., 2013).

RNA-seq reads can be linked to different genomic features, like genes and exons, using featureCounts to annotate them after alignment. FeatureCounts is a helpful tool that facilitates paired-end sequencing and makes read counting simpler (Liao et al., 2014).

**4. Differential expression using DESeq2**

Significant differences exist between genes and samples in discrete RNA-seq count data. *DESeq2* uses a statistical framework based on negative-binomial modeling and moderated estimates of dispersion and fold change to identify systematic differences between experimental conditions (Love et al., 2014). The use of biological duplicates and multiple-testing correction is essential because thousands of genes can be analyzed simultaneously.

**5. Pathway analysis and functional enrichment**

The three main categories of gene ontology are biological process, molecular function, and cellular component. These ontologies can be used to analyze gene lists according to common biological traits (Ashburner et al., 2000). g:Profiler, one of the most popular profiling tools, transforms gene IDs and helps identify categories with significant overrepresentation among the genes (Raudvere et al., 2019).

The KEGG database contains information about gene relationships, molecular activities, and pathways within biological systems (Kanehisa & Goto, 2000). KEGG overrepresentation analysis was performed on both upregulated and downregulated genes in this investigation. The nominal p-values could not be considered the final result of the adjusted p-values since the number of tests performed determines the significance of pathway enrichment.

#### RESEARCH OBJECTIVES

- The following goals guided the project's design:
- To gather and arrange publically accessible paired-end RNA-seq data that depicts normal tissue and breast tumors
- to assess the quality of sequencing both before and following read preprocessing.
- to keep paired-end reads while eliminating adapters and low-quality sequence segments.
- to use a splice-aware RNA-seq aligner to match the processed reads to the human reference genome.
- To use aligned reads to create gene-level read-count tables
- to use DESeq2 to find genes that are expressed differently in tumor and normal samples.
- to use PCA, a volcano plot, and a heatmap to display the differential-expression results.
- to determine which upregulated and downregulated genes belong to enriched Gene Ontology categories.
- to assess KEGG pathway overrepresentation following multiple-testing correction and suitable identifier conversion.
- To conservatively interpret the findings and record the entire process in report.

### METHODS

#### 1. Study design and data source

The publicly accessible RNA-seq dataset obtained from the NCBI Gene Expression Omnibus (GEO) was utilized for this project. GSE210787 was the selected series. *Twelve RNA-seq samples—six normal and six tumor*—are used in the experiment, which is categorized by the GEO database as an expression profiling experiment on Homo sapiens using Illumina NovaSeq 6000.

| **Parameter** | **Recorded information** |
| --- | --- |
| GEO accession | GSE210787 |
| Study title | Tumor-specific circRNAs elicit anti-tumor immune response via encoding cryptic peptides [RNA-Seq] |
| Organism | Homo sapiens |
| Platform | Illumina NovaSeq 6000 |
| Total samples | 12 |
| Tumor samples | 6 |
| Normal samples | 6 |
| Comparison | Tumor vs Normal |
| Read type | Paired-end |

Table 1. Dataset and experimental design**.**

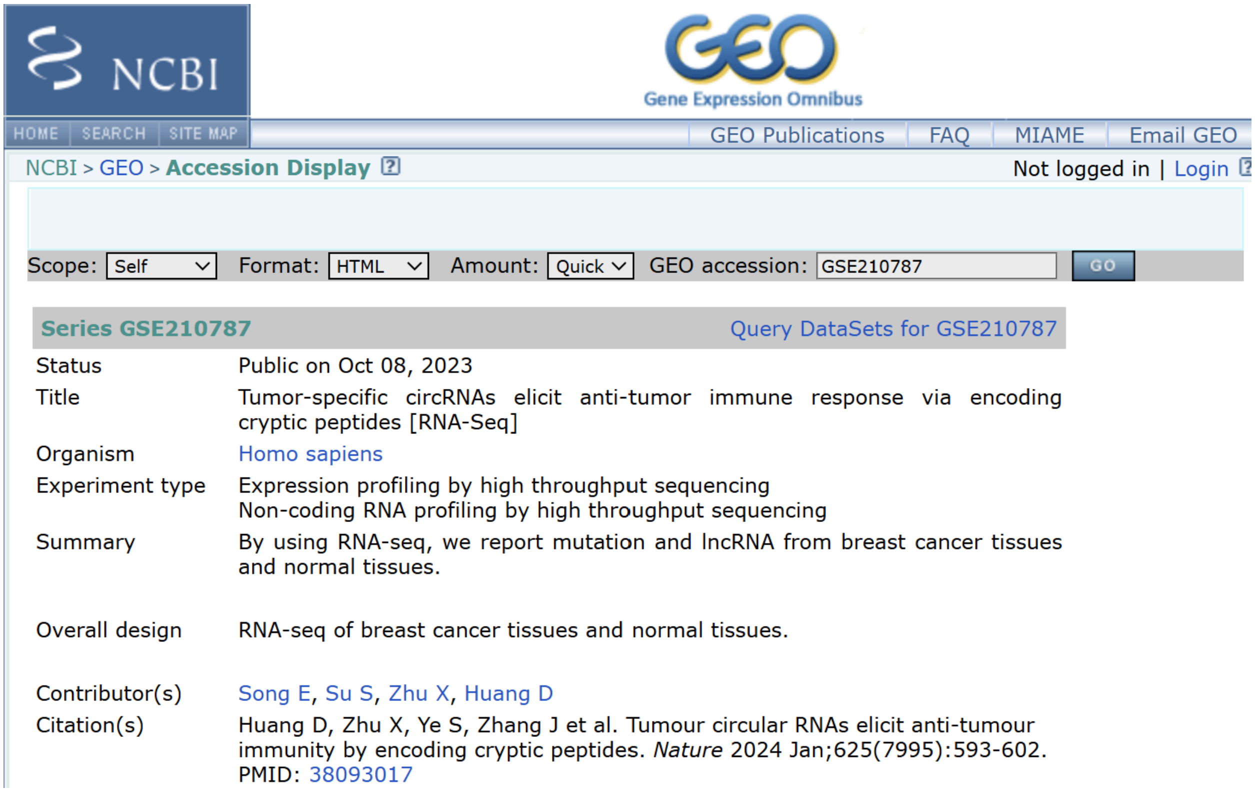

NCBI

GEO Gene Expression Omnibus

HOME | SEARCH | SITE MAP | GEO Publications | FAQ | MIAME | Email GEO

NCBI > GEO > Accession Display

Not logged in | Login

Scope: Self   Format: HTML   Amount: Quick   GEO accession: GSE210787   GO

| Series GSE210787 | Query DataSets for GSE210787 |
|---|---|
| Status | Public on Oct 08, 2023 |
| Title | Tumor-specific circRNAs elicit anti-tumor immune response via encoding cryptic peptides [RNA-Seq] |
| Organism | Homo sapiens |
| Experiment type | Expression profiling by high throughput sequencing<br>Non-coding RNA profiling by high throughput sequencing |
| Summary | By using RNA-seq, we report mutation and lncRNA from breast cancer tissues and normal tissues. |
| Overall design | RNA-seq of breast cancer tissues and normal tissues. |
| Contributor(s) | Song E, Su S, Zhu X, Huang D |
| Citation(s) | Huang D, Zhu X, Ye S, Zhang J et al. Tumour circular RNAs elicit anti-tumour immunity by encoding cryptic peptides. *Nature* 2024 Jan;625(7995):593-602. PMID: 38093017 |

Figure 1. Dataset/GEO.

#### 2. Sample and SRA run organization

| Sample | SRA run | Condition |
|---|---|---|
| T1 | SRR20982991 | Tumor |
| T2 | SRR20982994 | Tumor |
| T3 | SRR20982990 | Tumor |
| T4 | SRR20982989 | Tumor |
| T5 | SRR20982988 | Tumor |
| T6 | SRR20982987 | Tumor |
| N1 | SRR20982998 | Normal |
| N2 | SRR20982997 | Normal |
| N3 | SRR20982996 | Normal |

| | | |
|---|---|---|
| N4 | SRR20982995 | Normal |
| N5 | SRR20982993 | Normal |
| N6 | SRR20982992 | Normal |

Table 2. Sample-to-SRA run mapping maintained throughout the analysis.

**3. Computational environment and software**

| **Tool/resource** | **Version/release** | **Purpose** |
|---|---|---|
| SRA Toolkit | 3.2.1 | Raw-data retrieval |
| FastQC | 0.12.1 | Initial and post-trimming quality control |
| fastp | 1.0.1 | Adapter and quality trimming |
| MultiQC | 1.21 | Aggregation of QC reports |
| STAR | 2.7.11b+galaxy1 | Splice-aware RNA-seq alignment on Galaxy |
| featureCounts | 2.1.1+galaxy1 | Gene-level read counting |
| DESeq2 | 2.11.40.8+galaxy4 | Differential expression analysis |
| g:Profiler GOSt | 0.1.7+galaxy11 | GO enrichment |
| KEGG ORA Galaxy tool | Project run | KEGG over-representation analysis |
| GENCODE | Release 50 | Human gene annotation |
| Reference | GRCh38.p14 / hg38 | Human reference genome |

Table 3. Software, versions and purposes recorded.

**4. Reference genome and annotation**

The *GENCODE Release 50 annotation* and the human *GRCh38.p14* reference genome were utilized. The primary assembly FASTA file *GRCh38.primary_assembly.genome.fa* and gencode.v50.annotation were used in the project.GTF. Prior to alignment and counting, the annotation was examined and chromosome naming was verified for compatibility. A human reference annotation release linked to GRCh38.p14 is called GENCODE Release 50 (GENCODE, 2026).

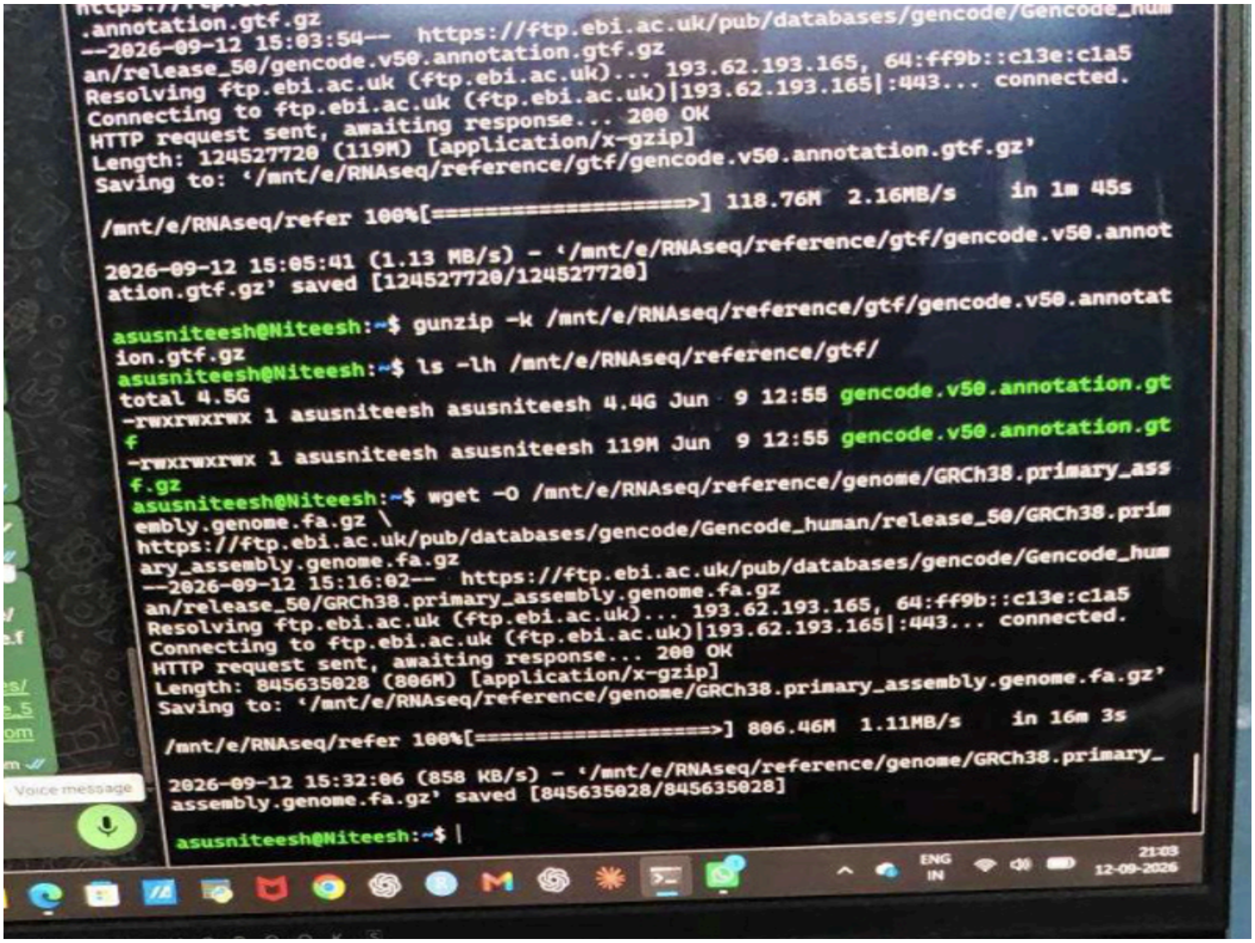


Figure 2. Reference Genome and Annotation Download.

**5. Raw data retrieval**

After being recovered, the twelve SRA runs were arranged as paired-end FASTQ files. Each run was kept connected to its intended tumor or normal condition by maintaining the sample metadata. The raw FASTQ directory was cleaned to free up storage space following verification and the successful creation of trimmed data.

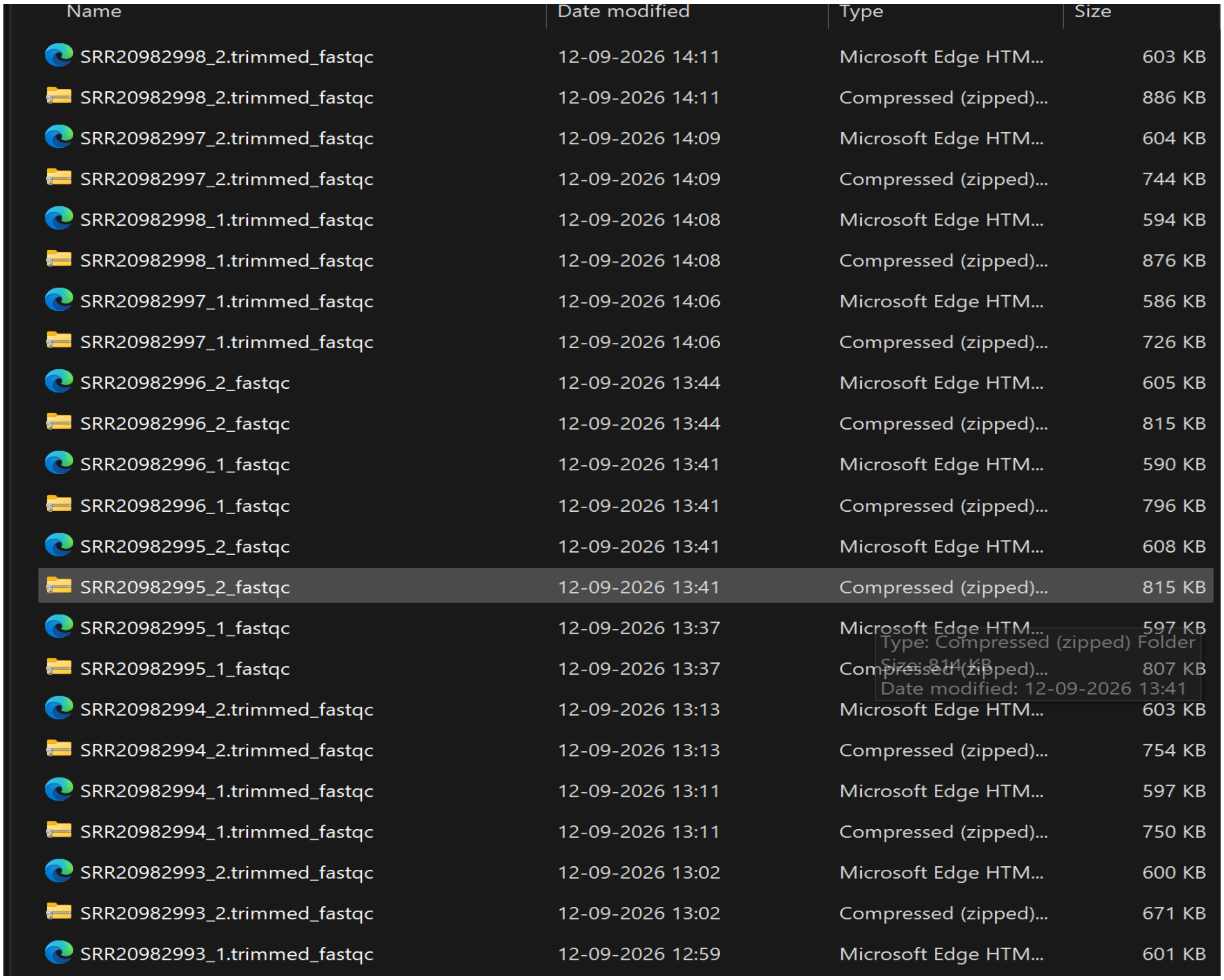


Figure 3. RNA-Seq Dataset and Trimmed Reads.

#### 6. Initial quality control

The raw reads were examined using FastQC. The project log shows that the visible samples have a GC content of roughly 49–52%, generally acceptable per-sequence quality, low levels of overrepresented sequences, and acceptable adapter content following trimming. Since duplication can happen in RNA-seq libraries as a result of transcript abundance and library composition, higher sequence duplication—roughly 50–60%—was noted rather than automatically treated as a sequencing failure in many samples. In the context of RNA-seq/priming bias, per-base sequence-content warnings were also observed and taken into consideration.

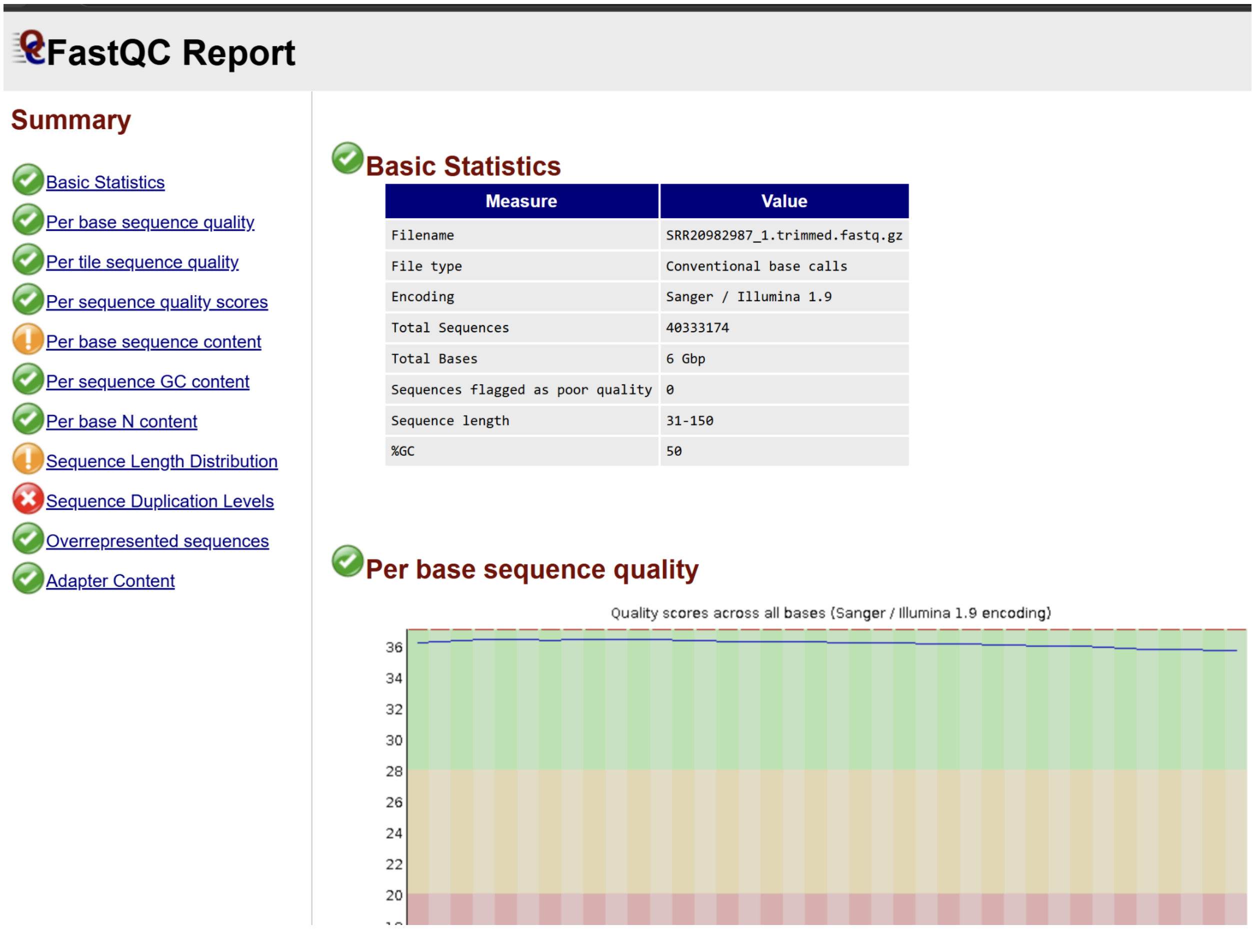


Figure 4. FastQC Quality Control Report.

#### 7. Read trimming and post-trimming QC

Fastp version 1.0.1 was used to preprocess paired-end sequencing reads. The purpose of this step was to remove low-quality bases and sequencing adapters from the sequences before mapping.The twelve samples have all been successfully preprocessed, preserving both read mates.

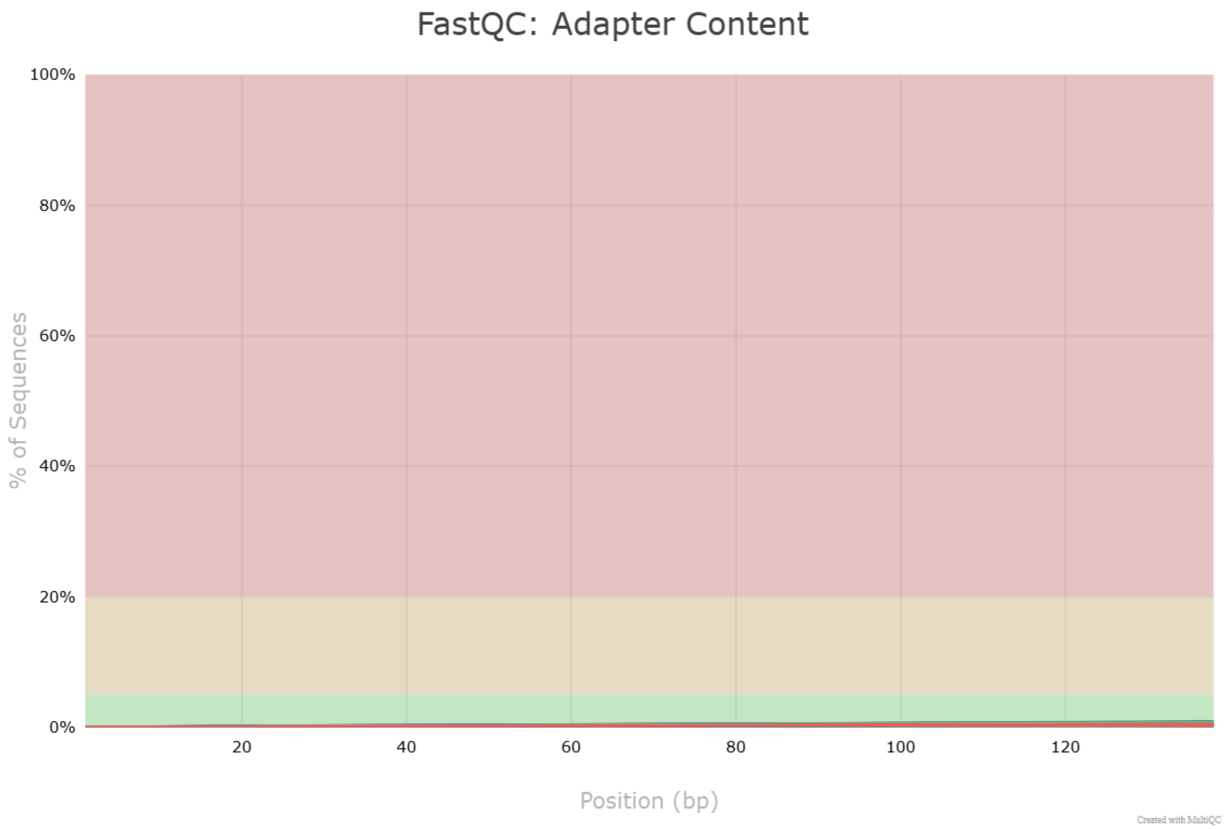


Figure 5. FastQC Analysis of Trimmed Reads.

**8. STAR alignment**

In WSL/Ubuntu, the project first tried local STAR genome indexing. Under the local resources available, the human index generation process repeatedly stalled during the junction-insertion stage. On the mentor's advice, the workflow was transferred to Galaxy instead of altering the intended aligner. A paired-end collection comprising the twelve sample pairs, the integrated human hg38 reference, and the GENCODE v50 GTF was used to run STAR 2.7.11b+galaxy1 in

Galaxy. Twelve BAM outputs were produced with success.

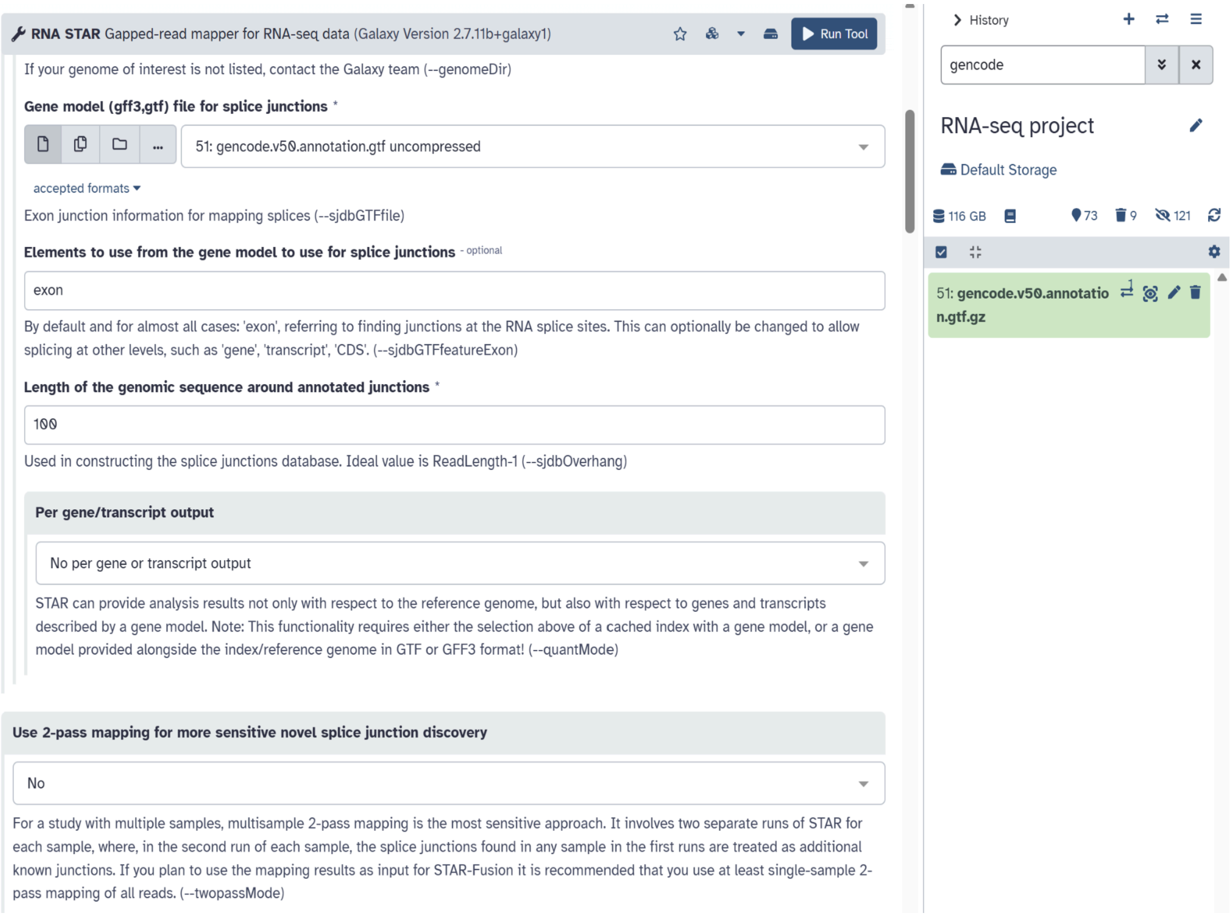


Figure 6. STAR Alignment Configuration.

#### 9. Gene-level quantification with featureCounts

The aligned data was subjected to featureCounts version 2.1.1+galaxy1. For gene/meta-feature-level counting, the project made use of exon features and the gene_id attribute. Reads with paired ends were tallied as fragments.The gene-count tables that were produced served as DESeq2's input.

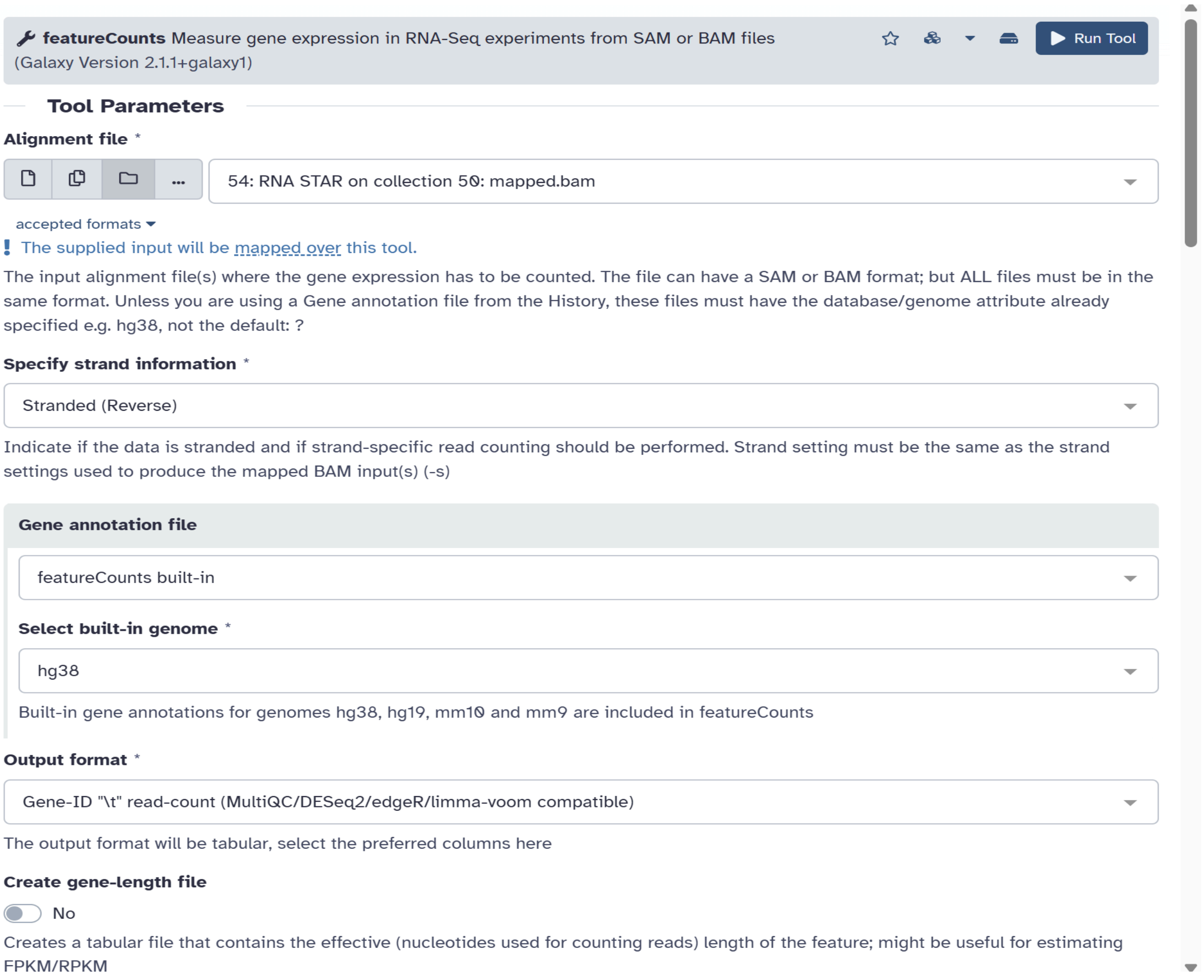


Figure 7. featureCounts Configuration.

#### 10. Differential expression with DESeq2

Six tumor samples and six normal samples were compared using DESeq2 version 2.11.40.8+galaxy4. Tumor versus Normal was the definition of the condition factor. The entire DESeq2 result was kept as Galaxy dataset #145. The study employed both an absolute log2 fold-change threshold of at least 1 and an adjusted p-value threshold of less than 0.05.

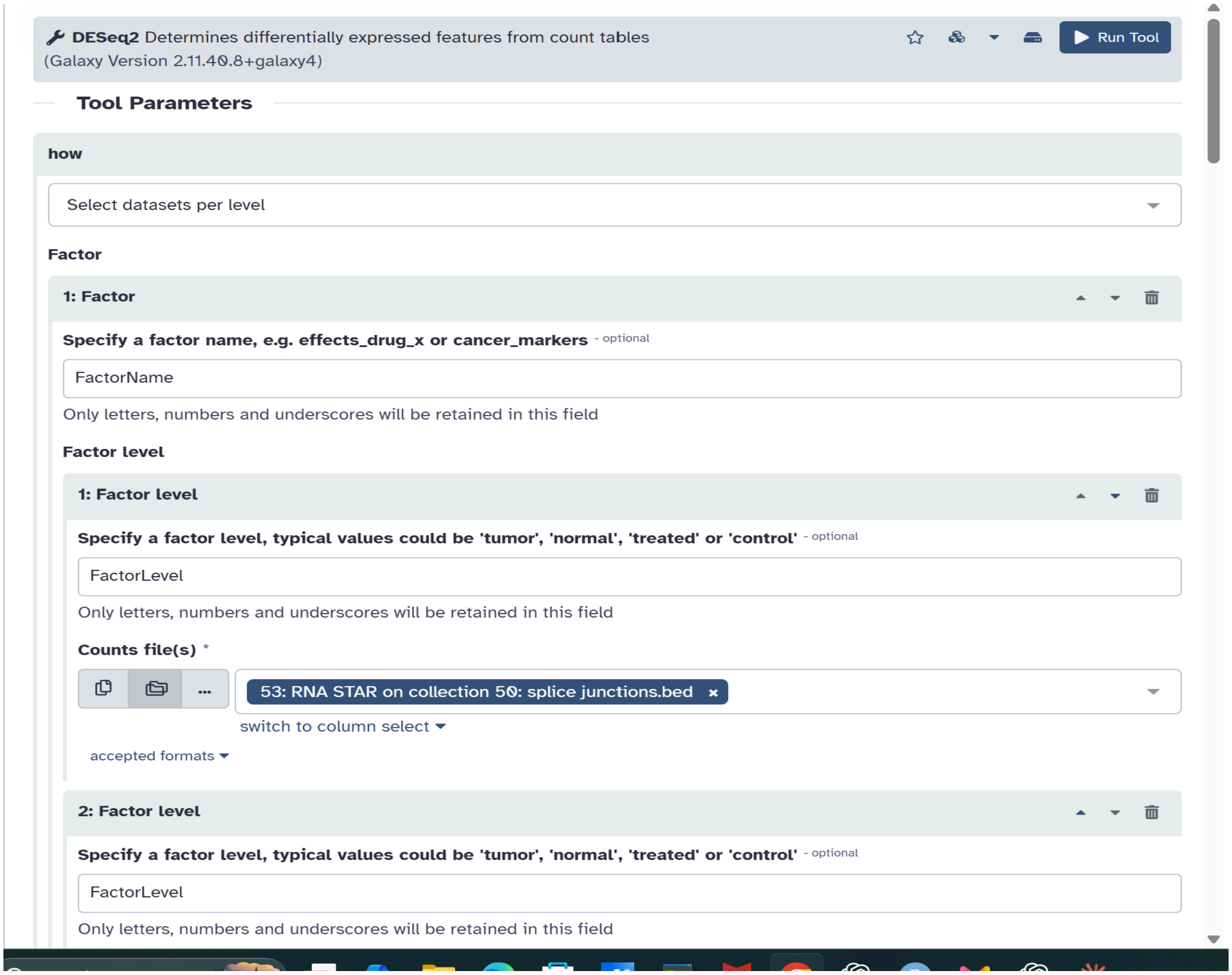


Figure 8. DESeq2 Parameters.

| Filtering criterion | Threshold |
|---|---|
| Adjusted p-value (padj) | < 0.05 |
| Absolute log2 fold change | ≥ 1 |

Table 4. Differential-expression selection criteria.

#### 11. Visualization

To see statistical significance and effect size, a volcano plot was created using the entire DESeq2 result (#145). The 120 chosen DEGs' VST-normalized expression values were used to create a heatmap. All twelve samples were included in the heatmap, which visualized expression patterns using sample clustering.

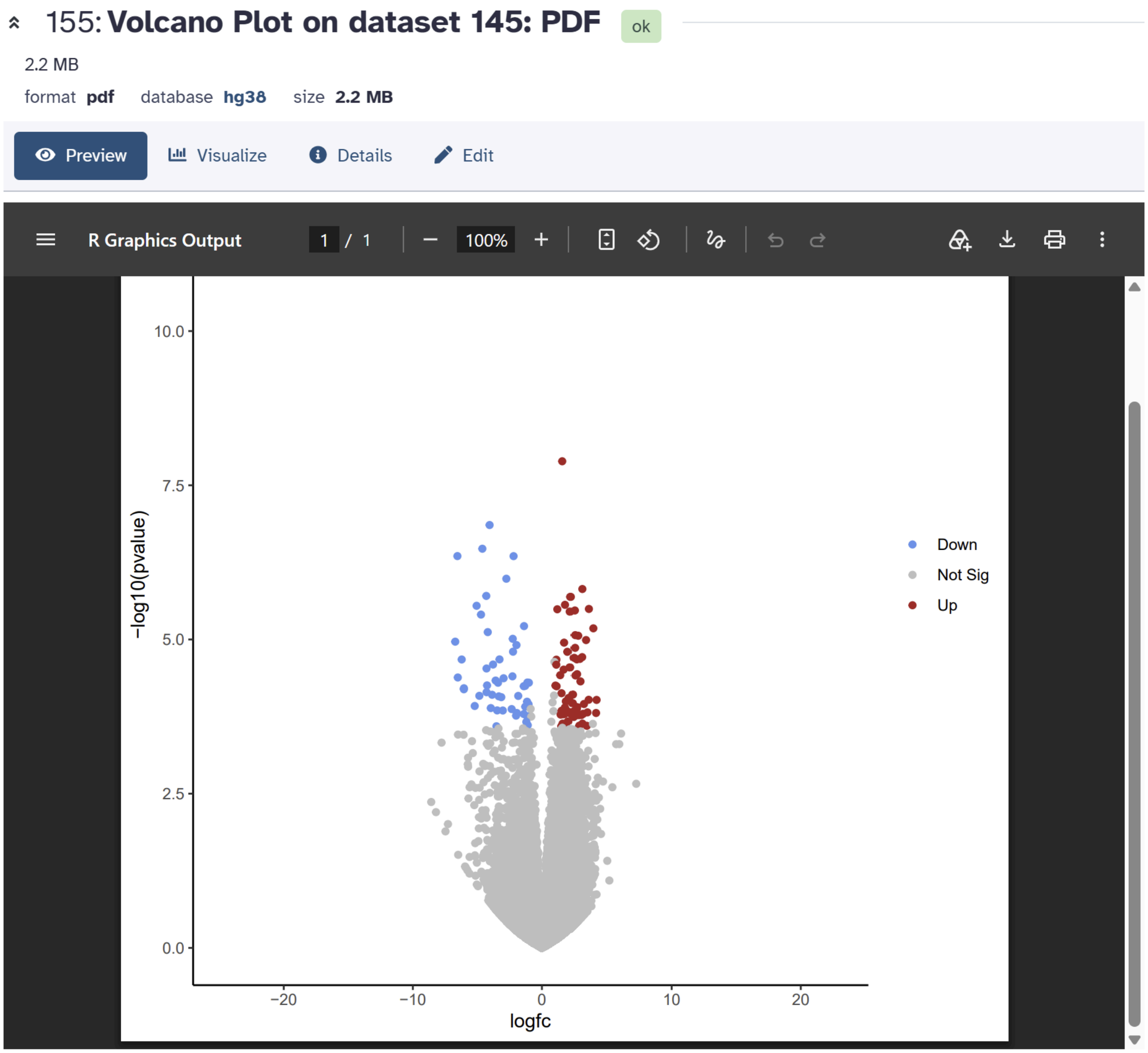


Figure 9. Volcano Plot of Differentially Expressed Genes.

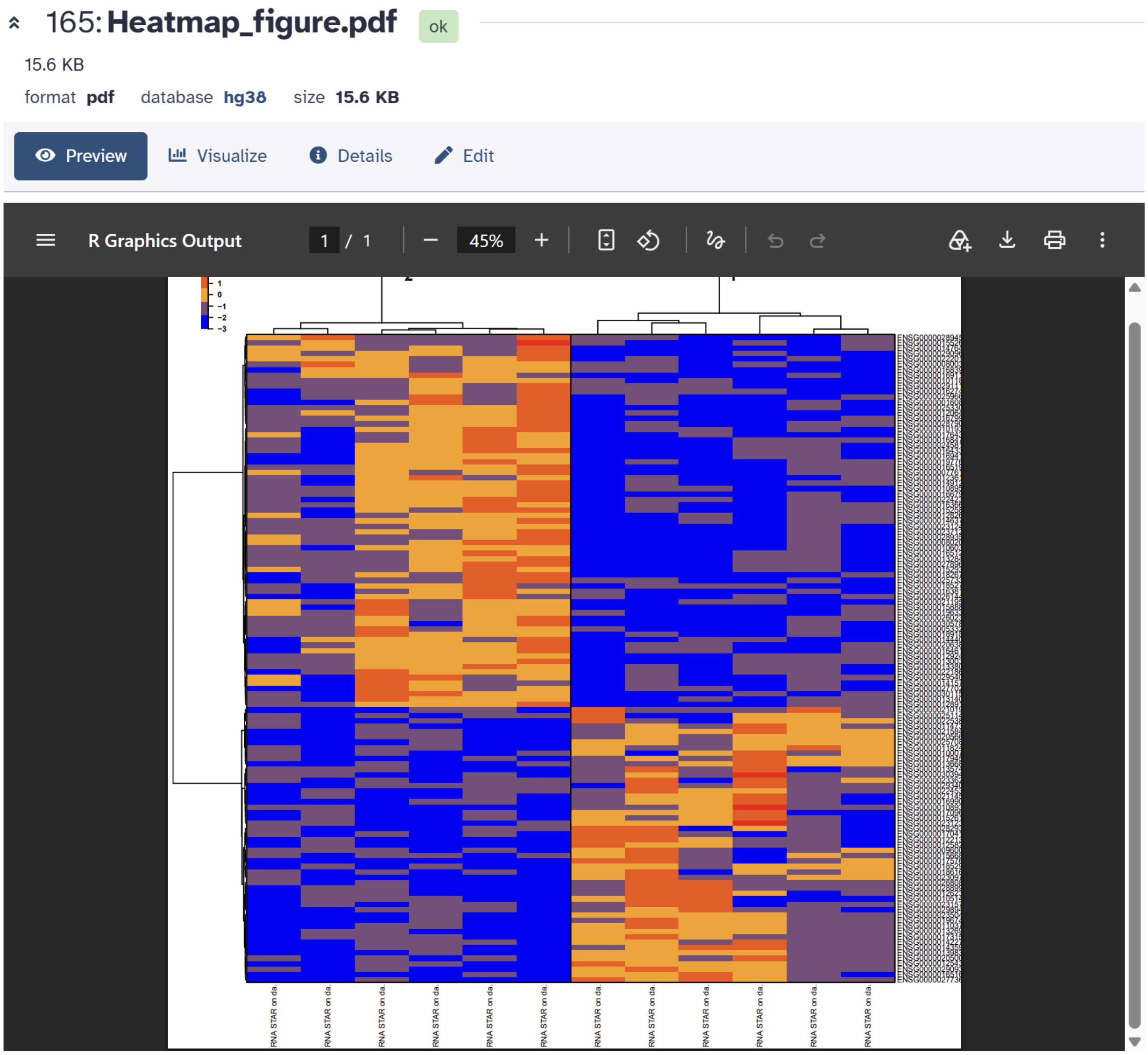


Figure 10. Heatmap Analysis Results.

#### 12. Gene Ontology enrichment

GO enrichment was performed separately for upregulated and downregulated gene lists using *g:Profiler GOSt*. The analysis considered Biological Process, Molecular Function, and Cellular Component categories. Ensembl version suffixes were removed before enrichment to improve identifier compatibility.

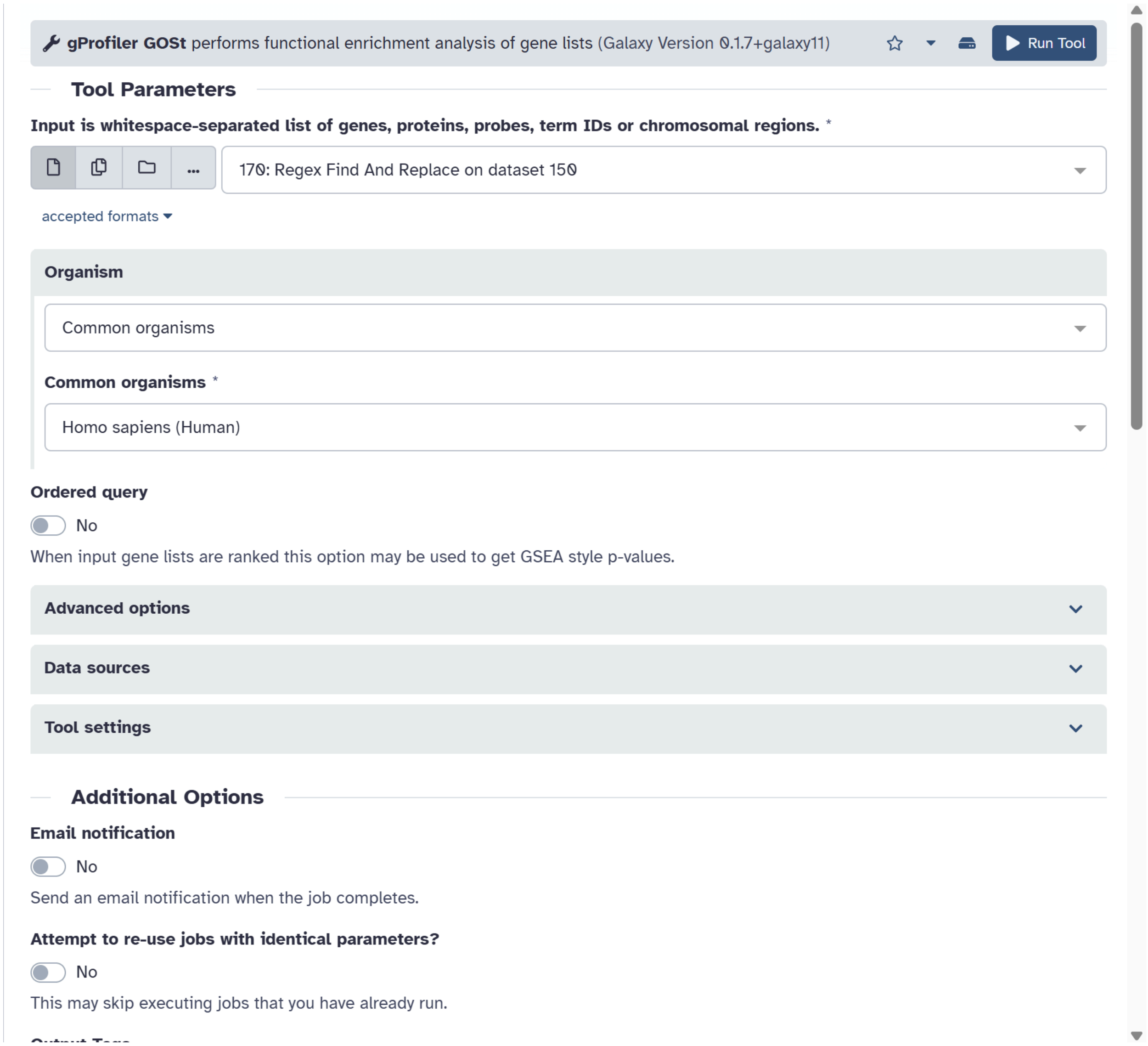


Figure 11. GO Functional Enrichment Analysis.

#### 13. KEGG pathway analysis

KEGG over-representation analysis was performed separately for the upregulated and downregulated gene sets. Ensembl identifiers were converted to Entrez identifiers before analysis, and unmapped identifiers were removed. The human KEGG organism code was used. The project log records that repeated g:Profiler attempts did not produce the expected KEGG terms, so the KEGG analysis was completed with a dedicated *KEGG ORA tool*.

#### RESULTS & DISCUSSION

##### 1. Data retrieval and sample organization

The final analysis comprised twelve RNA-seq samples in paired-end mode: six for tumors and six for normal. The association of the biological samples' names with the corresponding SRA runs was kept in all steps of the pipeline, which was crucial as the differential expression modeling required an accurate pairing of each count matrix to its condition.

##### 2. Quality-control findings

During the quality control step, it became apparent that the sequencing data could be progressed beyond the pre-processing phase. The measured GC percentages were quite narrow, lying around 49-52%, and per-sequence quality appeared to be fine in all examined QC reports. Adapter content, after the trimming, appeared to be acceptable. Sequences overrepresented in the examined data set were less than 1%.

The most important finding during the quality control process was the duplication of sequence. It was high in a number of the examined samples, but it was not taken by itself as a sign of failure of the experiment. In bulk RNA-seq, many sequences can duplicate naturally, thus duplications should always be considered in light of the biological background and other QC modules. Similarly, per-base sequence content warnings were noted but not automatically considered to be a failure of RNA-seq library preparation.

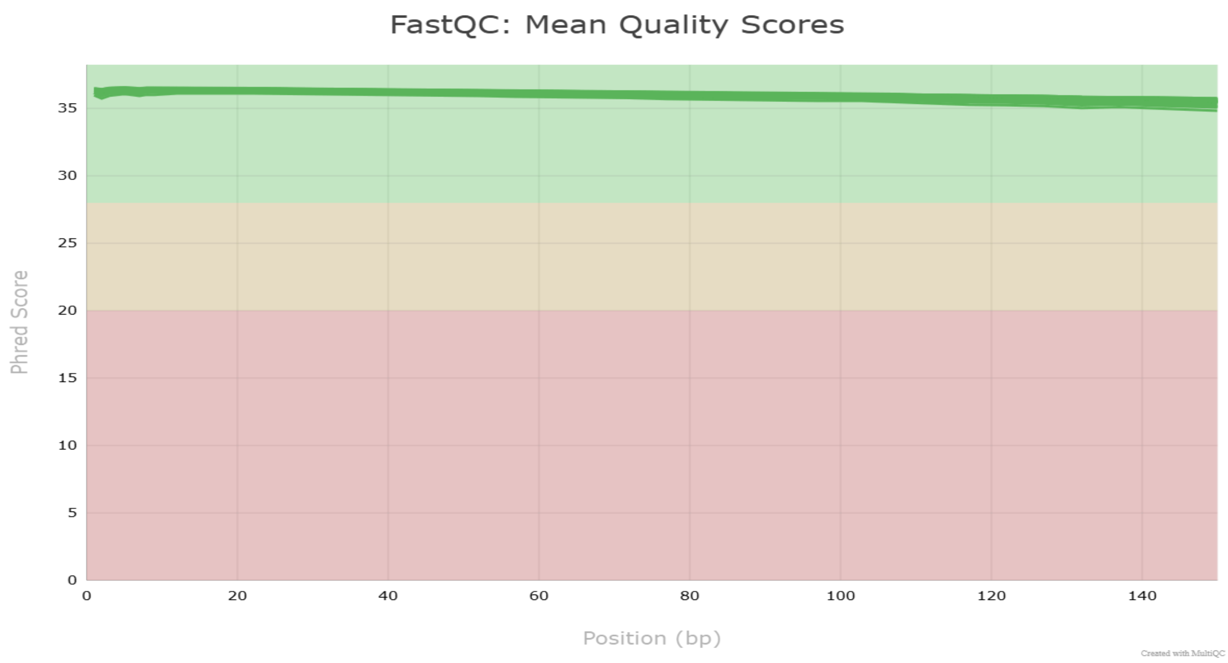


.

Figure 12. Per Base Sequence Quality

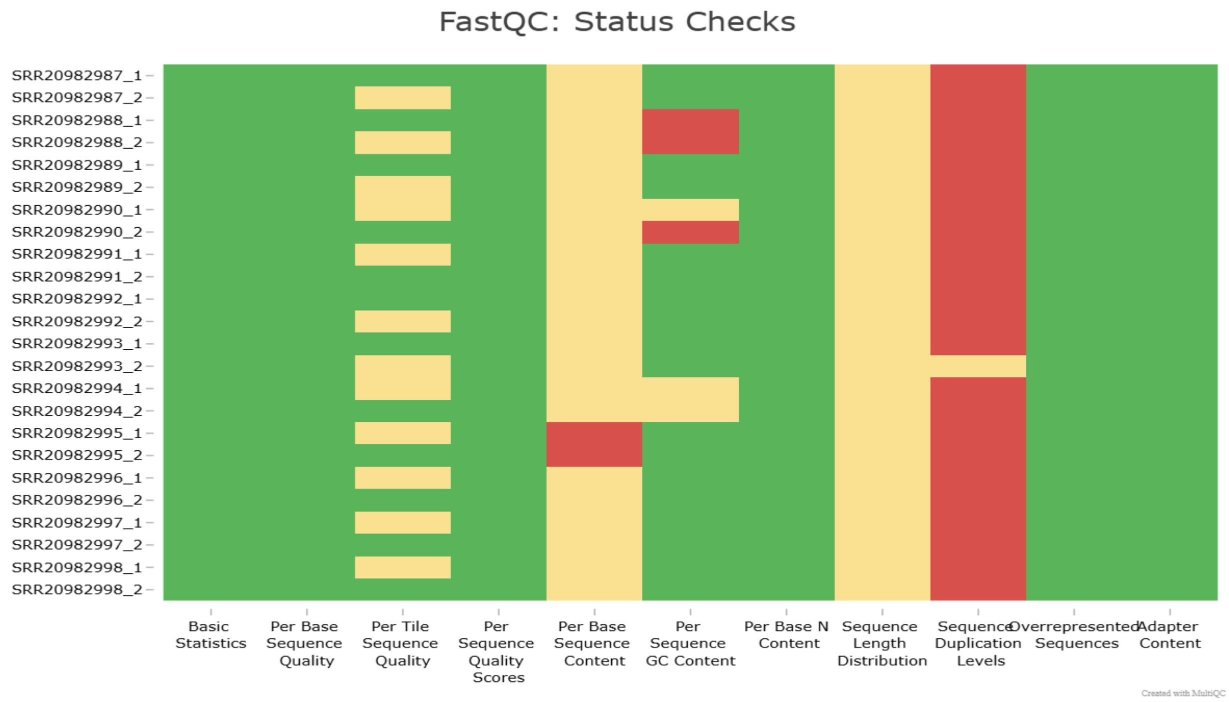


FastQC Status-Check Heatmap

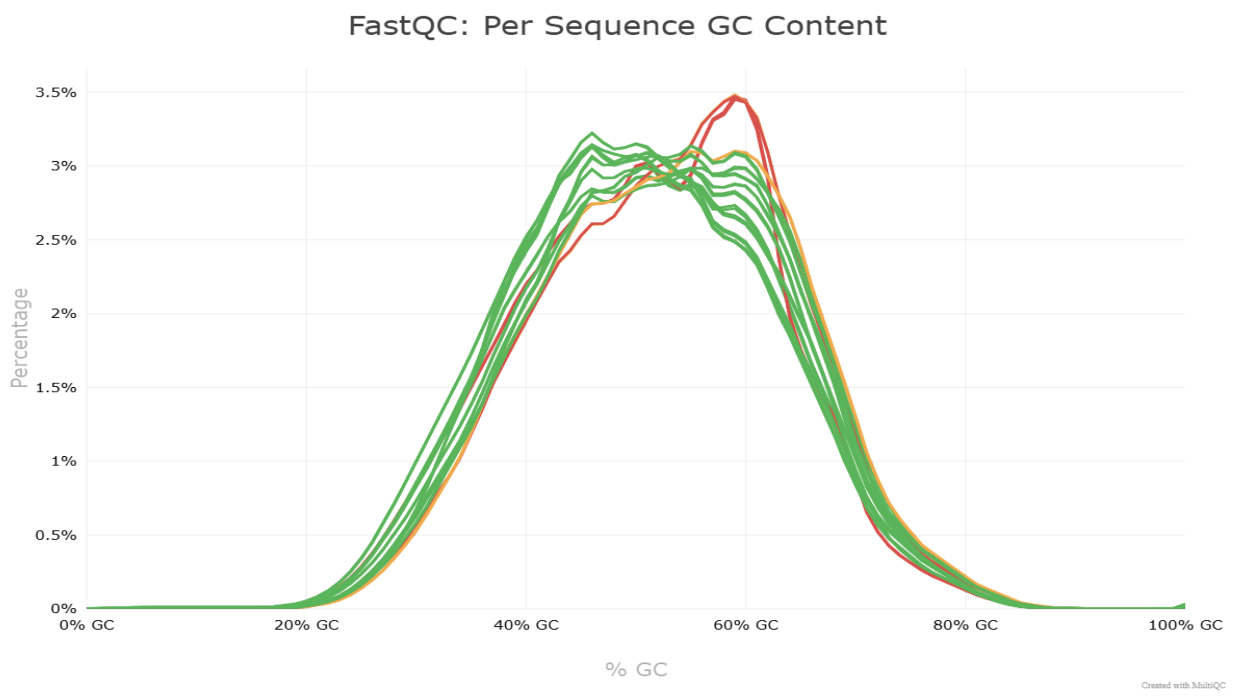


Per-sequence GC Content

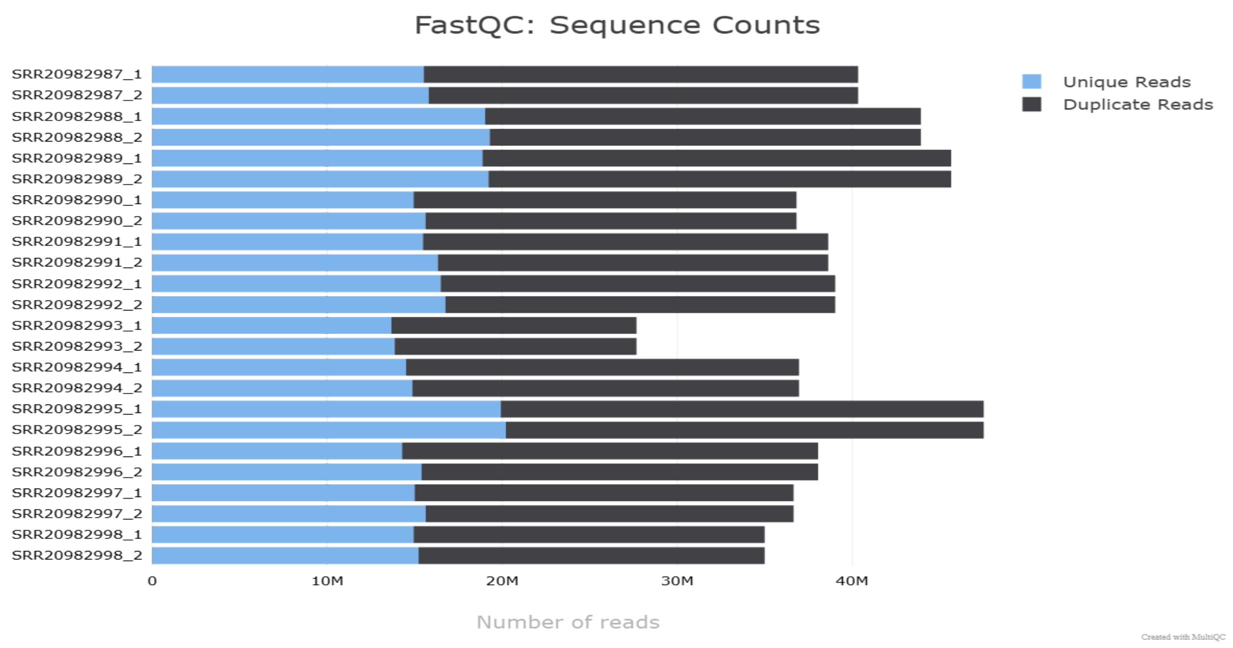


Sequence counts

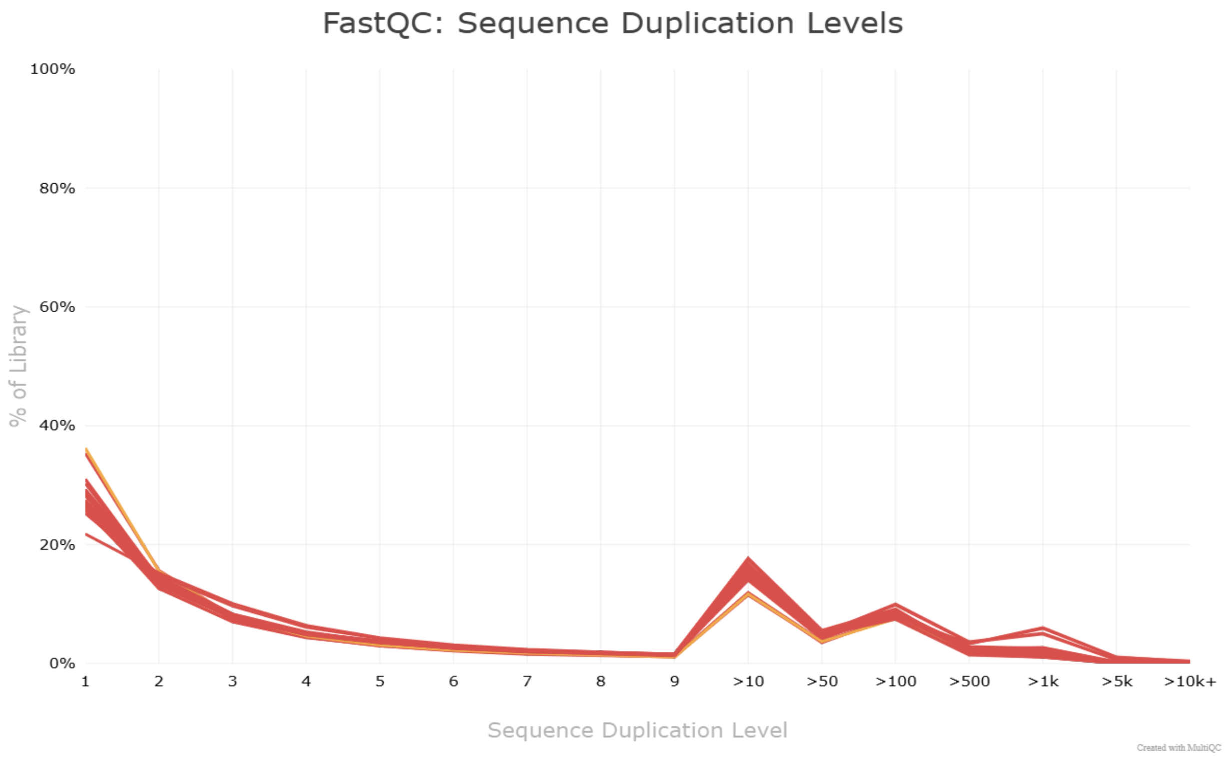


Sequence duplication levels.

| QC observation | Project finding | Interpretation used |
|---|---|---|
| GC content | GC-content distributions are concentrated mainly around 45–65%, with peaks varying among samples. | GC profiles are broadly similar, with some variation between samples. |
| Per-sequence quality | Mean quality scores remain approximately around Phred 33 in the displayed plot. | Overall per-base quality appears high in the displayed summary. |
| Adapter content | Adapter-content status is predominantly green in the MultiQC heatmap. | No widespread adapter-content warning is evident in the displayed summary. |
| Overrepresented sequences | Status is predominantly green in the MultiQC heatmap. | No widespread overrepresented-sequence warning is evident in the displayed summary. |
| Sequence duplication | Duplication-level distributions vary across samples; the exact percentages are not confirmed by these screenshots. | Duplication should be interpreted in the context of RNA-seq libraries and other QC metrics. |
| Per-base sequence content | Warnings are visible for some samples in the MultiQC heatmap. | The warnings should be considered alongside the remaining QC results |

Table 5. Summary of quality-control observations.

##### 3. STAR alignment and BAM generation

The mentor's intended alignment strategy was not abandoned in order to solve the local STAR indexing issue. The analysis could proceed with STAR and the chosen hg38/GENCODE reference configuration by shifting the computationally intensive alignment stage to Galaxy. *Twelve BAM outputs* were successfully generated, providing the aligned-read files needed for gene counting downstream.

#### 91: RNA STAR on dataset 51: log

ok

36 lines

format **txt** database **hg38** size **2 KB**

Preview Visualize Details Edit

```
                                 Started job on |       Sep 21 05:03:15
                             Started mapping on |       Sep 21 06:12:02
                                    Finished on |       Sep 21 06:28:07
       Mapping speed, Million of reads per hour |       163.84

                          Number of input reads |       43918386
                      Average input read length |       299
                                    UNIQUE READS:
                   Uniquely mapped reads number |       39510555
                        Uniquely mapped reads % |       89.96%
                          Average mapped length |       297.39
                       Number of splices: Total |       36526050
            Number of splices: Annotated (sjdb) |       35593027
                       Number of splices: GT/AG |       36149888
                       Number of splices: GC/AG |       235089
                       Number of splices: AT/AC |       26568
               Number of splices: Non-canonical |       114505
                      Mismatch rate per base, % |       0.32%
                         Deletion rate per base |       0.01%
                        Deletion average length |       1.92
                        Insertion rate per base |       0.01%
                       Insertion average length |       1.60
                             MULTI-MAPPING READS:
        Number of reads mapped to multiple loci |       3122956
             % of reads mapped to multiple loci |       7.11%
        Number of reads mapped to too many loci |       29528
             % of reads mapped to too many loci |       0.07%
                                  UNMAPPED READS:
  Number of reads unmapped: too many mismatches |       0
       % of reads unmapped: too many mismatches |       0.00%
            Number of reads unmapped: too short |       1201509
                 % of reads unmapped: too short |       2.74%
                Number of reads unmapped: other |       53838
                     % of reads unmapped: other |       0.12%
                                  CHIMERIC READS:
                       Number of chimeric reads |       0
                            % of chimeric reads |       0.00%
```

Figure 13. STAR Alignment BAM Output.

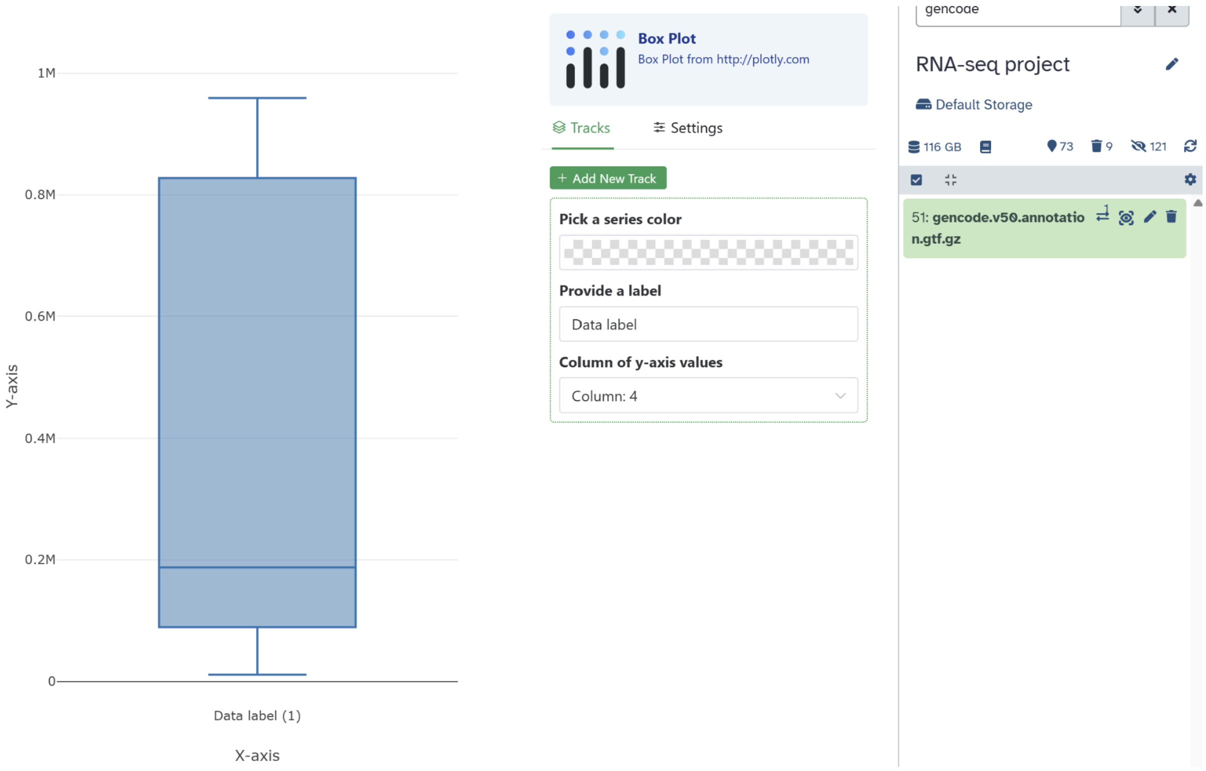


#### 4. Gene-level quantification

For each of the twelve samples, featureCounts generated gene-level count tables. The count data used in DESeq2 was consistently constructed using exon features and the gene_id attribute. The two mates from the same fragment were not treated as separate biological observations by counting paired-end reads as fragments.

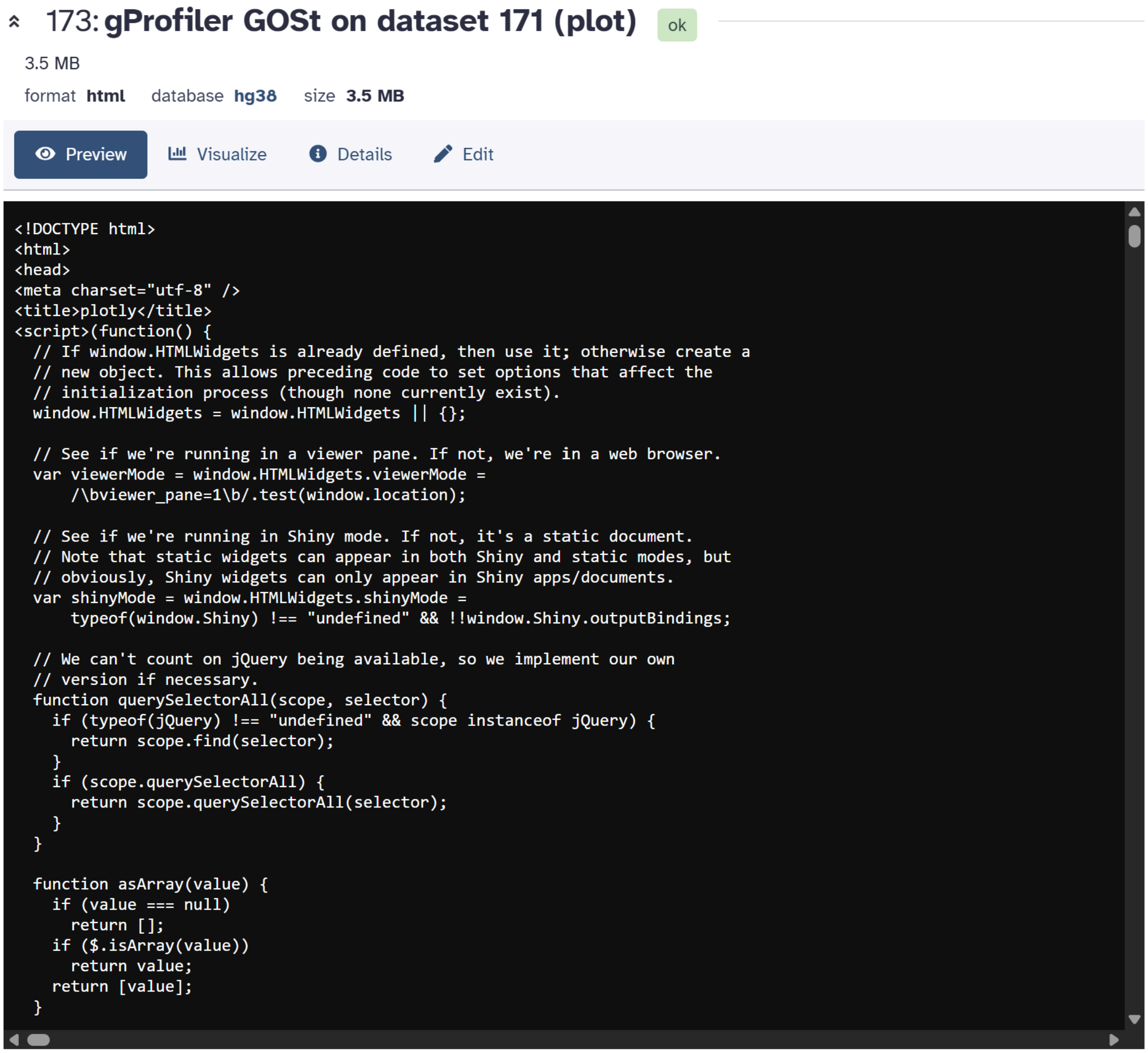


Figure 14. featureCounts Gene Count Results.

#### 5. Differential expression results

Identified *120 genes that met the predetermined thresholds* were found by the DESeq2 comparison. Of these, *51 were found to be downregulated and 69 to be upregulated* in the tumor group. As a result, the outcome demonstrated a discernible transcriptional difference between the two experimental settings.

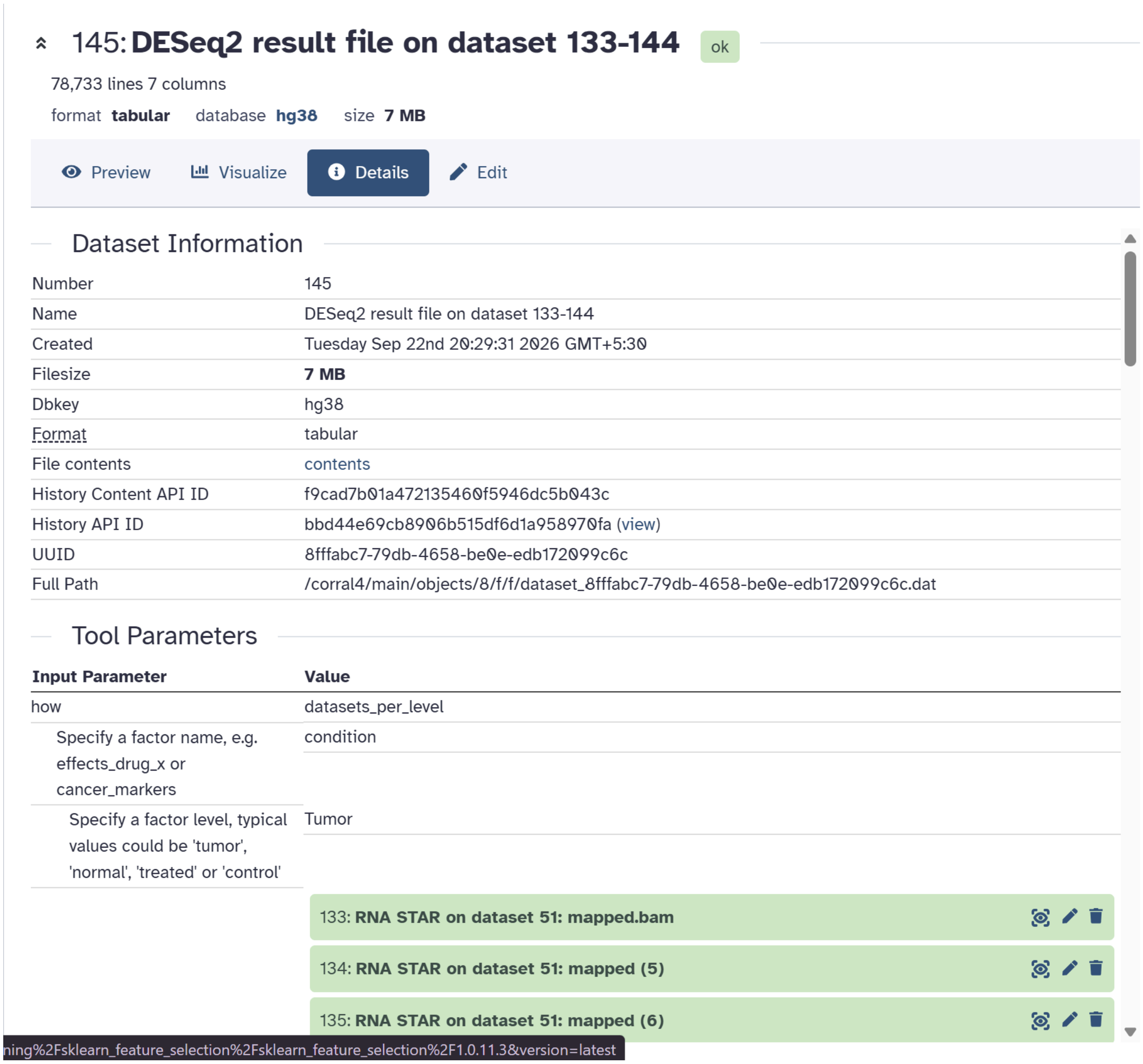


Figure 15. DESeq2 Differential Expression Analysis.

#### 146: DESeq2 plots on dataset 133-144

ok

10.9 MB

format **pdf** database **hg38** size **10.9 MB**

Preview Visualize Details Edit

R Graphics Output 1 / 5 100%

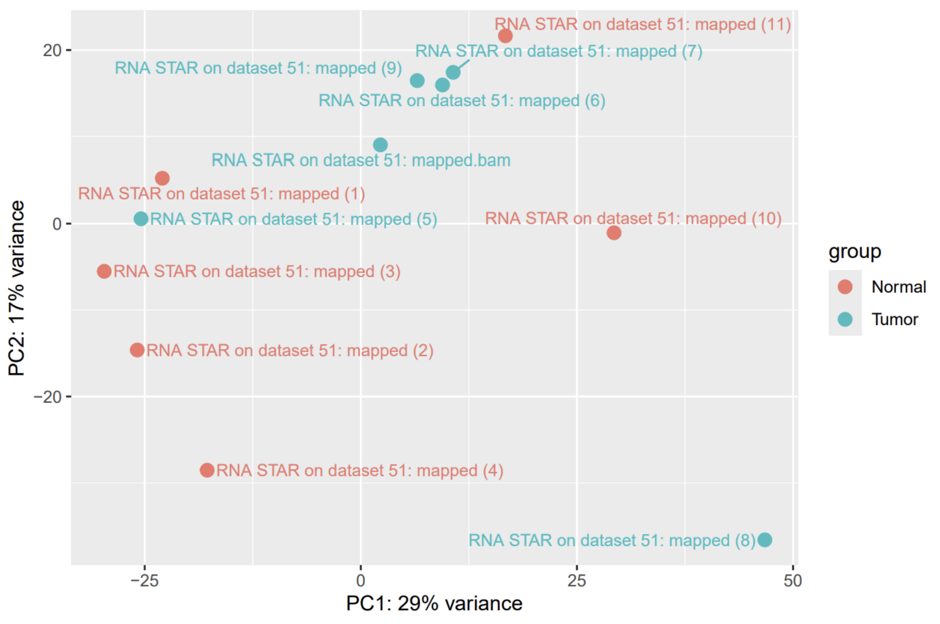

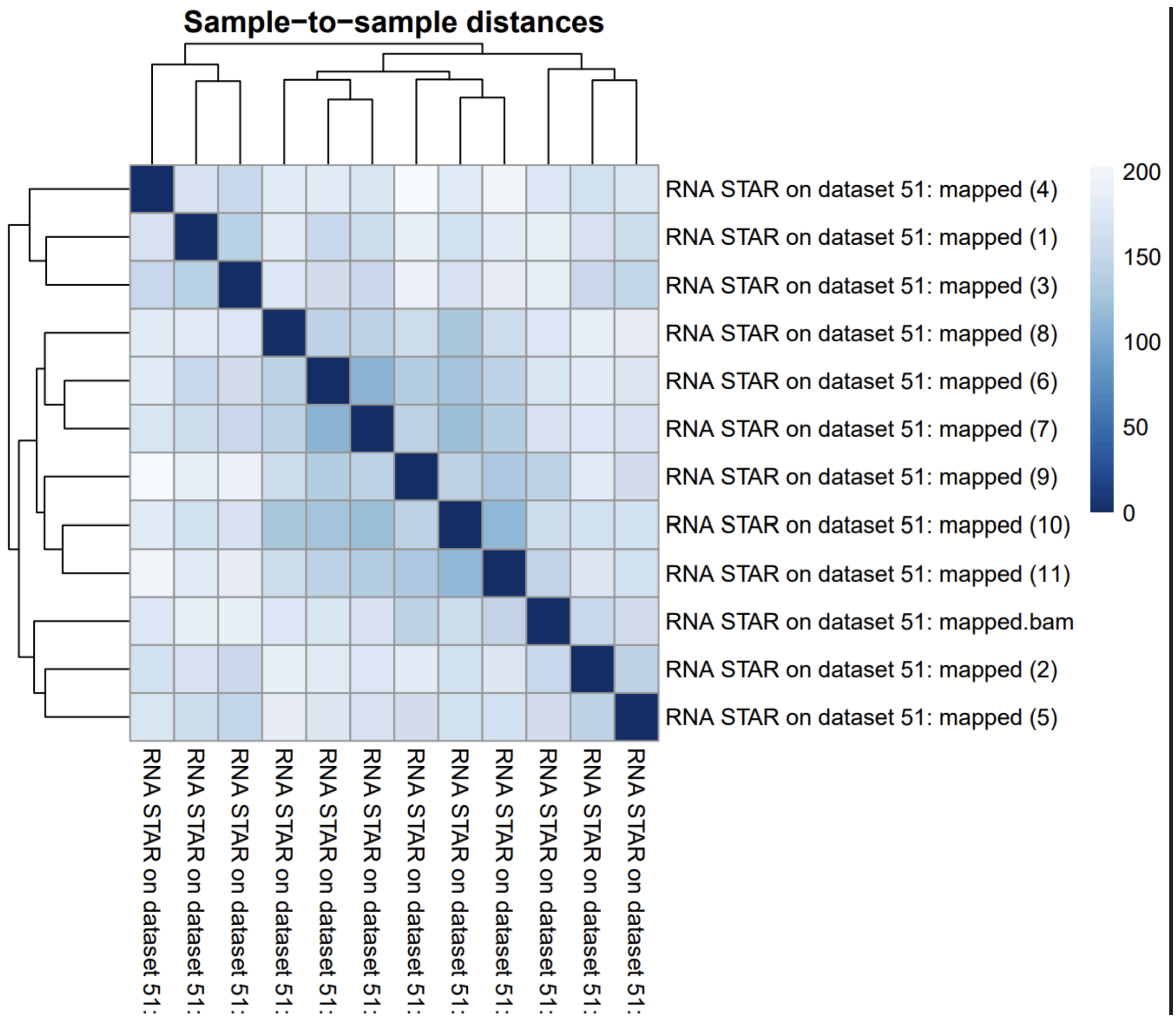
Sample−to−sample distances
RNA STAR on dataset 51: mapped (4)
RNA STAR on dataset 51: mapped (1)
RNA STAR on dataset 51: mapped (3)
RNA STAR on dataset 51: mapped (8)
RNA STAR on dataset 51: mapped (6)
RNA STAR on dataset 51: mapped (7)
RNA STAR on dataset 51: mapped (9)
RNA STAR on dataset 51: mapped (10)
RNA STAR on dataset 51: mapped (11)
RNA STAR on dataset 51: mapped.bam
RNA STAR on dataset 51: mapped (2)
RNA STAR on dataset 51: mapped (5)
200
150
100
50
0
RNA STAR on dataset 51:
RNA STAR on dataset 51:
RNA STAR on dataset 51:
RNA STAR on dataset 51:
RNA STAR on dataset 51:
RNA STAR on dataset 51:
RNA STAR on dataset 51:
RNA STAR on dataset 51:
RNA STAR on dataset 51:
RNA STAR on dataset 51:
RNA STAR on dataset 51:
RNA STAR on dataset 51:

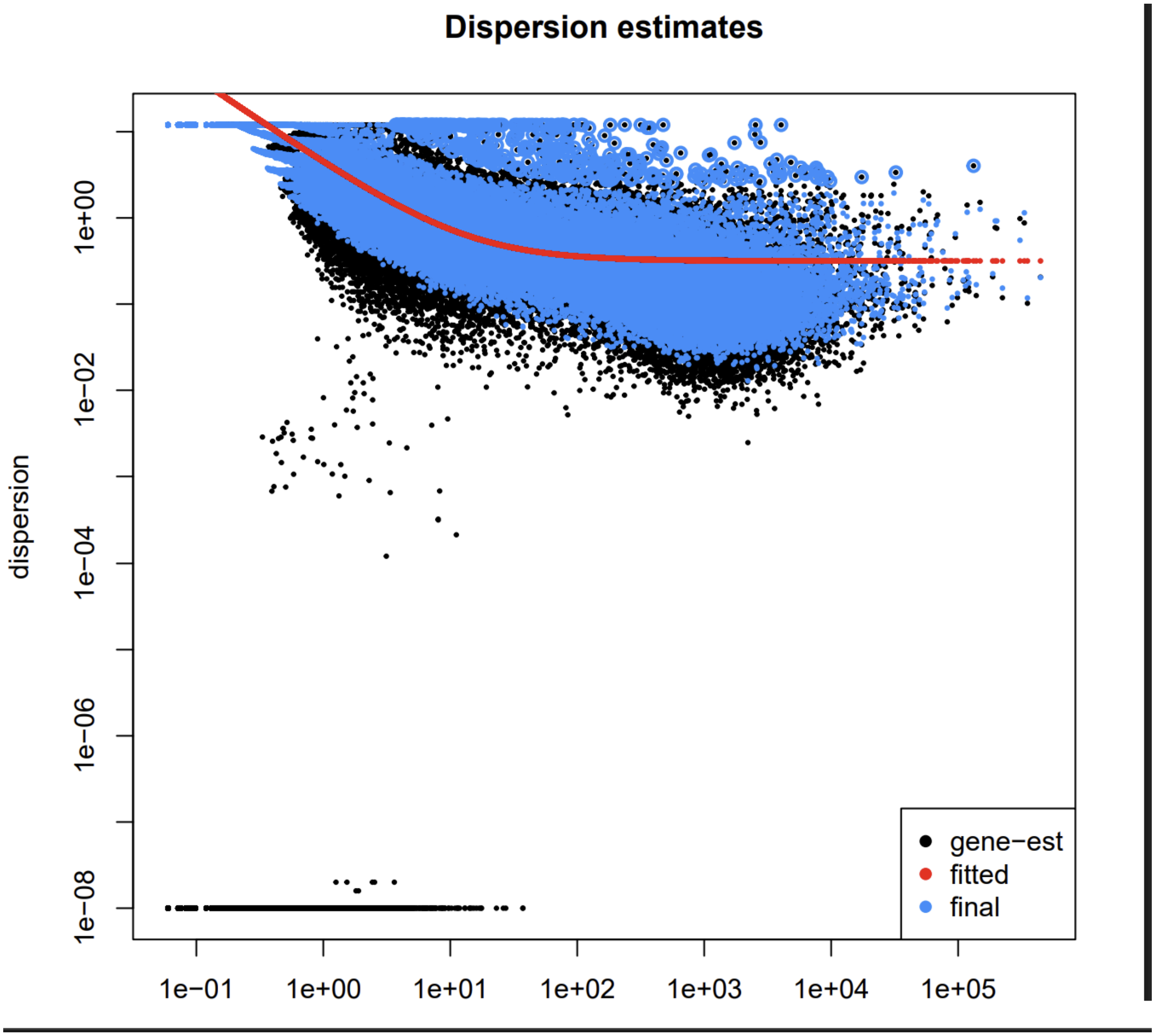

Dispersion estimates
dispersion
1e+00
1e−02
1e−04
1e−06
1e−08
1e−01
1e+00
1e+01
1e+02
1e+03
1e+04
1e+05
gene−est
fitted
final

**Histogram of p−values for condition: Tumor vs Normal**

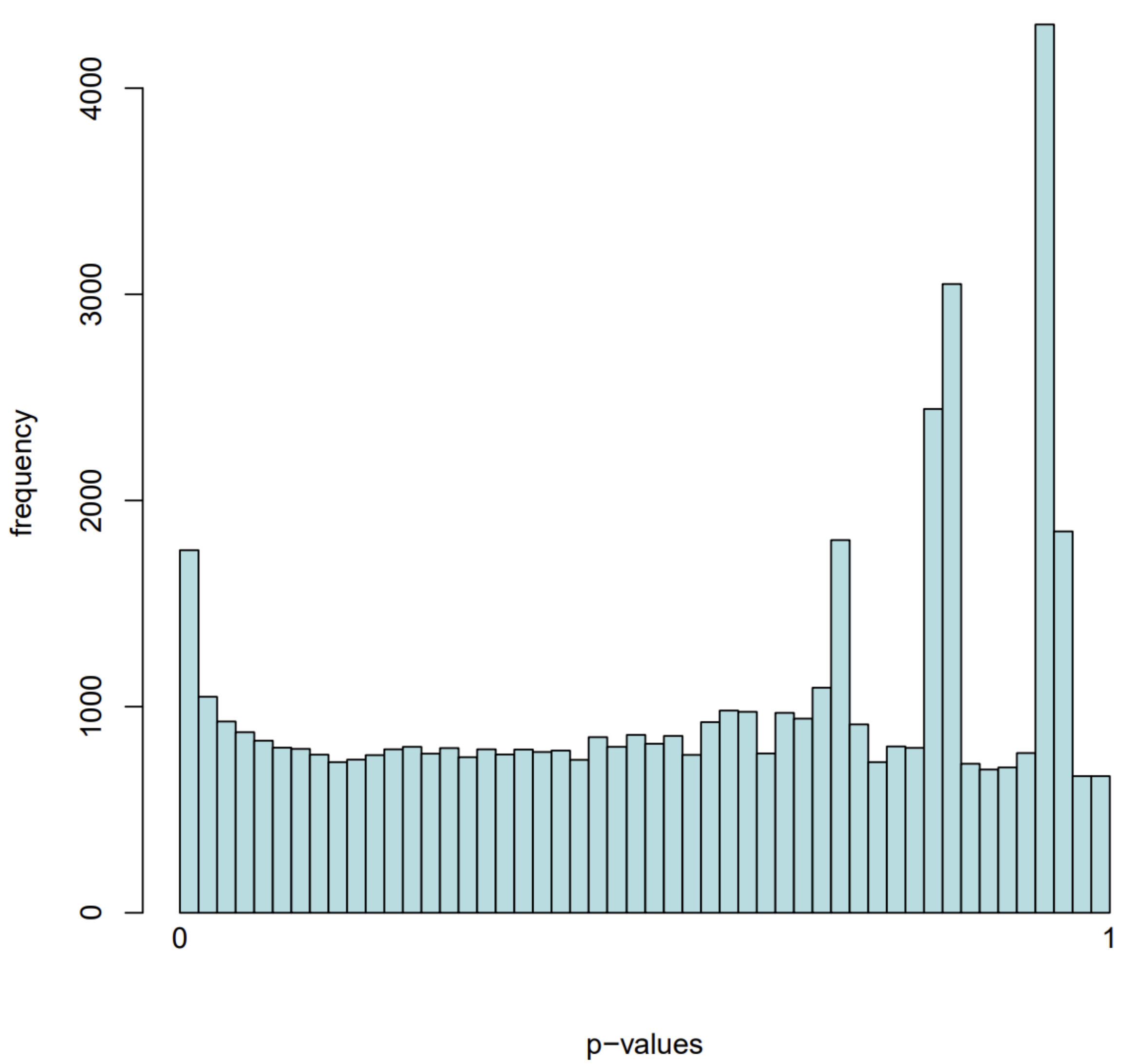

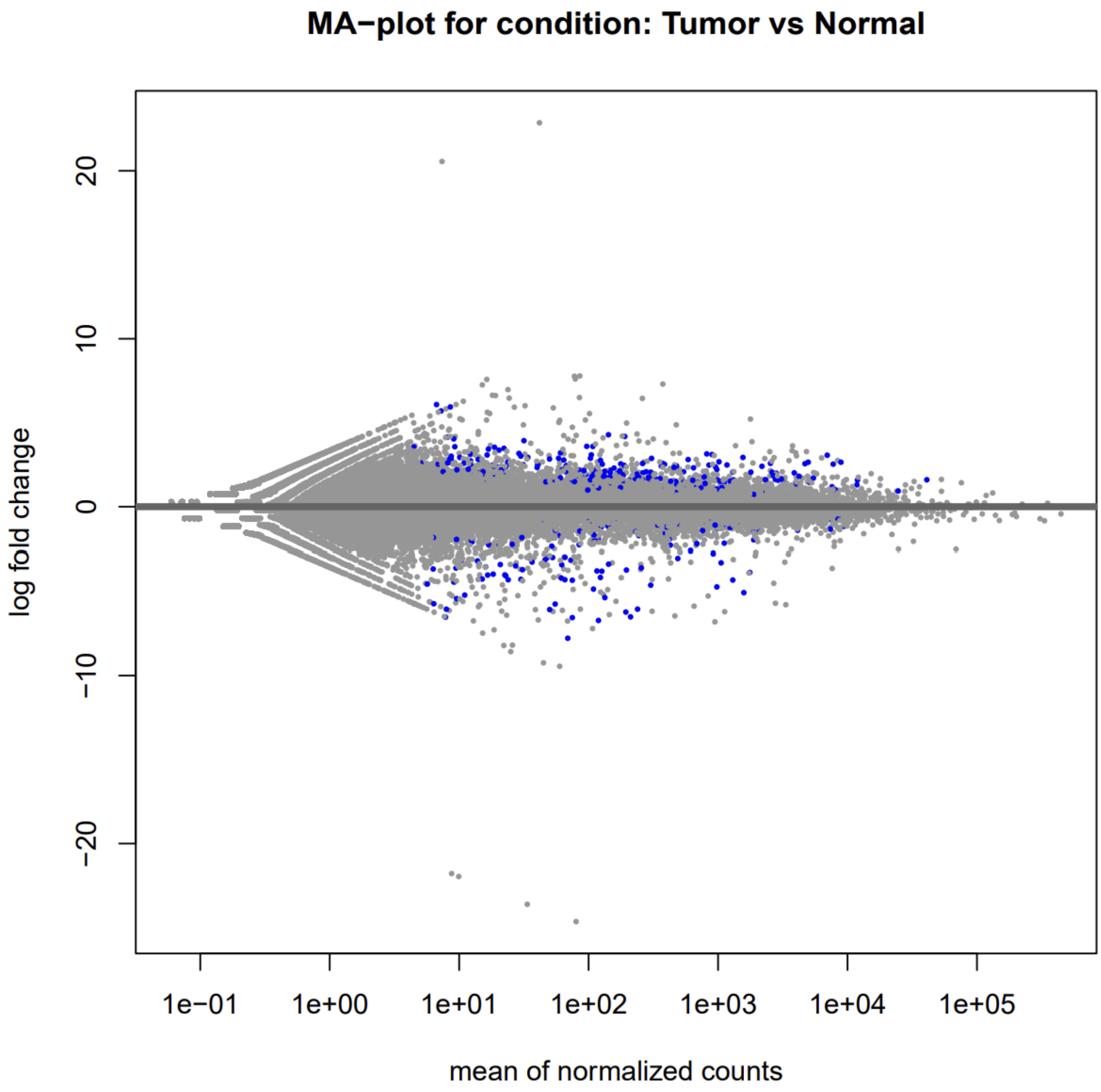


Figure 16.  DESeq2 PCA Plot.

| **Category** | **Number of genes** |
|---|---|
| Significant DEGs | 120 |
| Upregulated | 69 |
| Downregulated | 51 |

Table 6. Final differential-expression counts.

A condensed representation of the connection between effect size and statistical significance was offered by the volcano plot. The tumor and normal samples displayed different expression patterns in the heatmap based on the chosen DEGs. The observation that the chosen DEG set captures significant group-level transcriptional differences is supported by the sample clustering, which divides the two groups into major expression patterns.

**6. Gene Ontology enrichment**

The transition from gene lists to biological understanding was aided by enrichment analysis. Only one GO term from the list of *Cellular Components—GO:0071944*, *cell periphery*—turned out to be significant in terms of the up-regulated genes following functional enrichment. 29 genes were involved in the intersection, and the corresponding p-value was roughly 0.017585.

Cellular components located at or close to the cell's outer border are referred to as the "cell periphery." The significance of genes linked to this cellular component suggests some changes in cell-surface structures or cell-environment interactions when comparing tumors to normal. The findings of the enrichment analysis should indicate a functional relationship rather than a single mechanism.

With the selected enrichment settings, no GO terms among the down-regulated genes could be found. This negative result should be noted because it does not imply that down-regulated genes have no biological significance.

#### 182: **gProfiler GOSt on dataset 168** ok

1,932 lines 14 columns

format **tabular** database **hg38** size **276.4 KB**

Preview | Visualize | Details | Edit

| recall | term_id | source | term_name | effective_domain_size | source_order | parents |
|---|---|---|---|---|---|---|
| 855580412627 | GO:0048731 | GO:BP | system development | 20972 | 12138 | GO:0007275,GO:0048856 |
| 0.075 | GO:0001941 | GO:BP | postsynaptic membrane organization | 20972 | 540 | GO:0061024,GO:0099173 |
| 0.2 | GO:0097688 | GO:BP | glutamate receptor clustering | 20972 | 17491 | GO:0072578 |
| 0.2 | GO:0097113 | GO:BP | AMPA glutamate receptor clustering | 20972 | 17285 | GO:0001941,GO:0097688 |
| 81818181818182 | GO:1900449 | GO:BP | regulation of glutamate receptor signaling pathway | 20972 | 19408 | GO:0007215,GO:0009966 |
| 9913443830571 | GO:0044057 | GO:BP | regulation of system process | 20972 | 10196 | GO:0003008,GO:0051239 |
| 389830508475 | GO:0030334 | GO:BP | regulation of cell migration | 20972 | 6339 | GO:0016477,GO:2000145 |
| 3742828214647 | GO:0065008 | GO:BP | regulation of biological quality | 20972 | 14785 | GO:0065007 |
| 3846153846154 | GO:0021559 | GO:BP | trigeminal nerve development | 20972 | 5735 | GO:0021545 |
| 7121721587088 | GO:0048856 | GO:BP | anatomical structure development | 20972 | 12247 | GO:0032502 |
| 022696929239 | GO:0000165 | GO:BP | MAPK cascade | 20972 | 54 | GO:0141124 |

Figure 17. GO Enrichment Analysis Results.

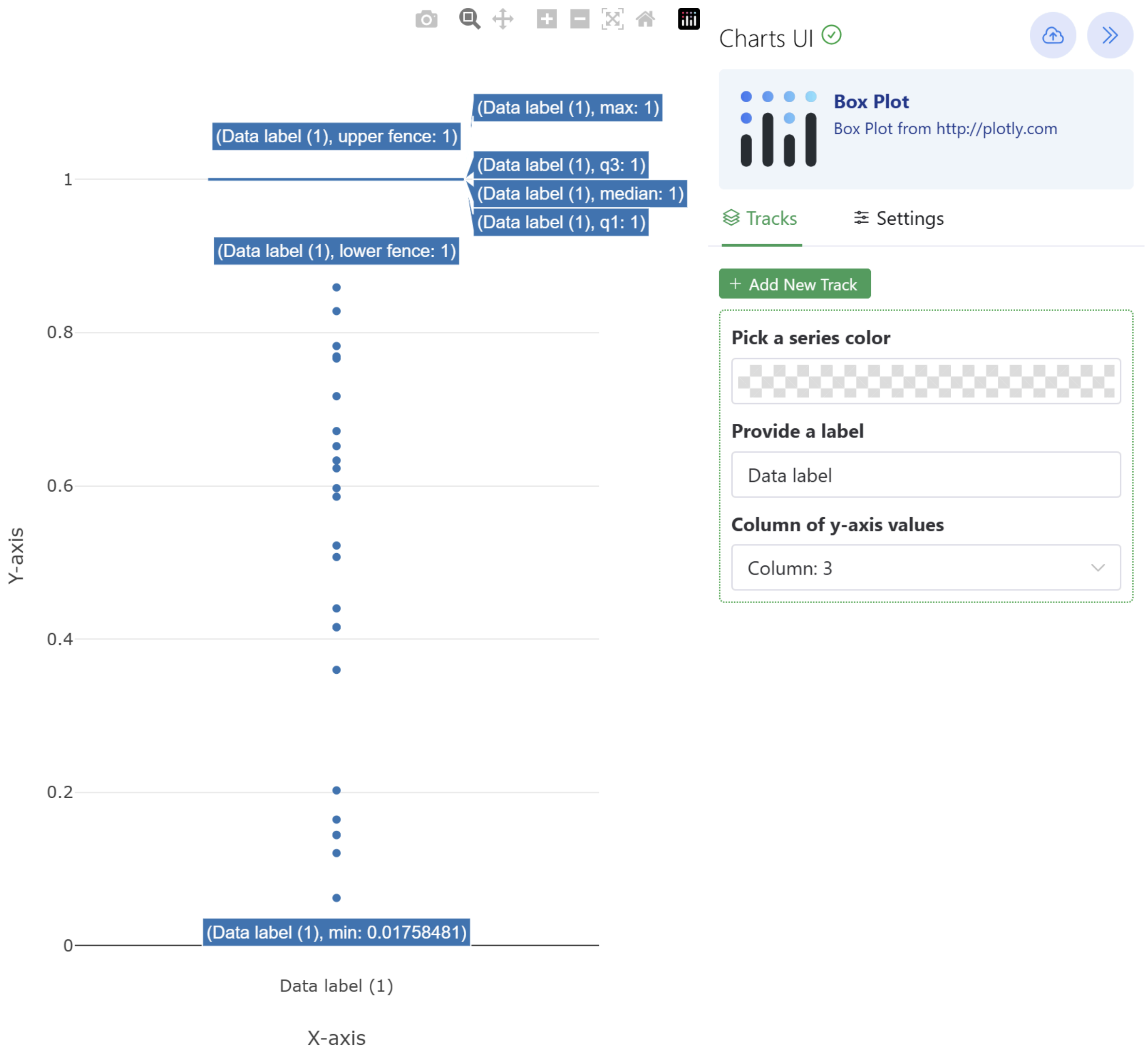


| Gene set | GO result | Recorded evidence |
|---|---|---|
| Upregulated | GO:0071944 — cell periphery (GO:CC) | p ≈ 0.017585; 29-gene intersection |
| Downregulated | No significant GO term | No category passed the selected significance criterion |

Table 7. GO enrichment summary.

#### 7. KEGG pathway enrichment

KEGG analysis was performed following the mapping of Ensembl identifiers to Entrez identifiers and the removal of unmapped entries. Several pathways showed nominal associations at $p < 0.05$ in the upregulated and downregulated gene sets, but none of them were statistically significant after multiple-testing correction.

For the upregulated gene set, the lowest a*djusted p-value was approximately 0.71393*, whereas the downregulated gene set had adjusted *p-values of 1.0.* Consequently, no KEGG pathway met the study's adjusted significance threshold.

Testing takes place concurrently. This distinction between enrichment in multiple pathways. Multiple-testing correction reduces the likelihood of interpreting pathways as significant simply because many statistical tests were performed.

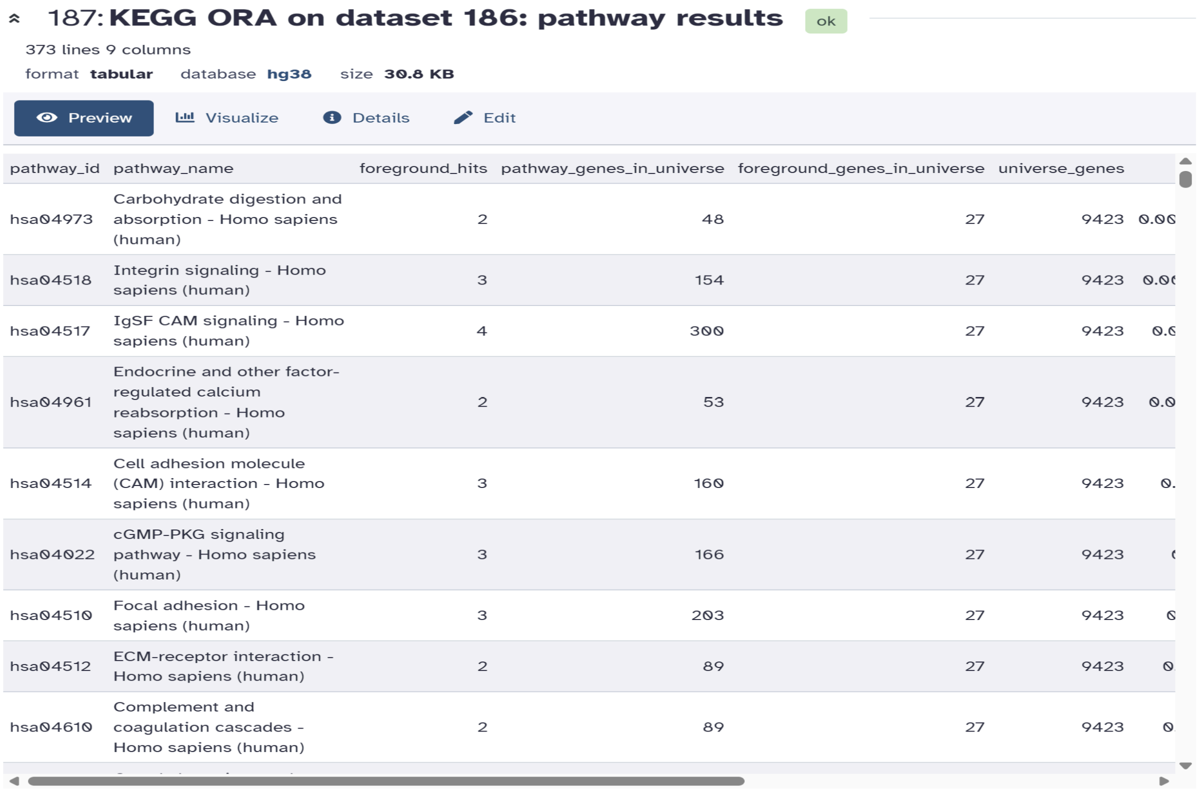

187: **KEGG ORA on dataset 186: pathway results** ok

373 lines 9 columns

format **tabular** database **hg38** size **30.8 KB**

Preview | Visualize | Details | Edit

| pathway_id | pathway_name | foreground_hits | pathway_genes_in_universe | foreground_genes_in_universe | universe_genes | |
|---|---|---|---|---|---|---|
| hsa04973 | Carbohydrate digestion and absorption - Homo sapiens (human) | 2 | 48 | 27 | 9423 | 0.00 |
| hsa04518 | Integrin signaling - Homo sapiens (human) | 3 | 154 | 27 | 9423 | 0.0 |
| hsa04517 | IgSF CAM signaling - Homo sapiens (human) | 4 | 300 | 27 | 9423 | 0.0 |
| hsa04961 | Endocrine and other factor-regulated calcium reabsorption - Homo sapiens (human) | 2 | 53 | 27 | 9423 | 0.0 |
| hsa04514 | Cell adhesion molecule (CAM) interaction - Homo sapiens (human) | 3 | 160 | 27 | 9423 | 0. |
| hsa04022 | cGMP-PKG signaling pathway - Homo sapiens (human) | 3 | 166 | 27 | 9423 | [illegible] |
| hsa04510 | Focal adhesion - Homo sapiens (human) | 3 | 203 | 27 | 9423 | 0 |
| hsa04512 | ECM-receptor interaction - Homo sapiens (human) | 2 | 89 | 27 | 9423 | 0 |
| hsa04610 | Complement and coagulation cascades - Homo sapiens (human) | 2 | 89 | 27 | 9423 | 0 |

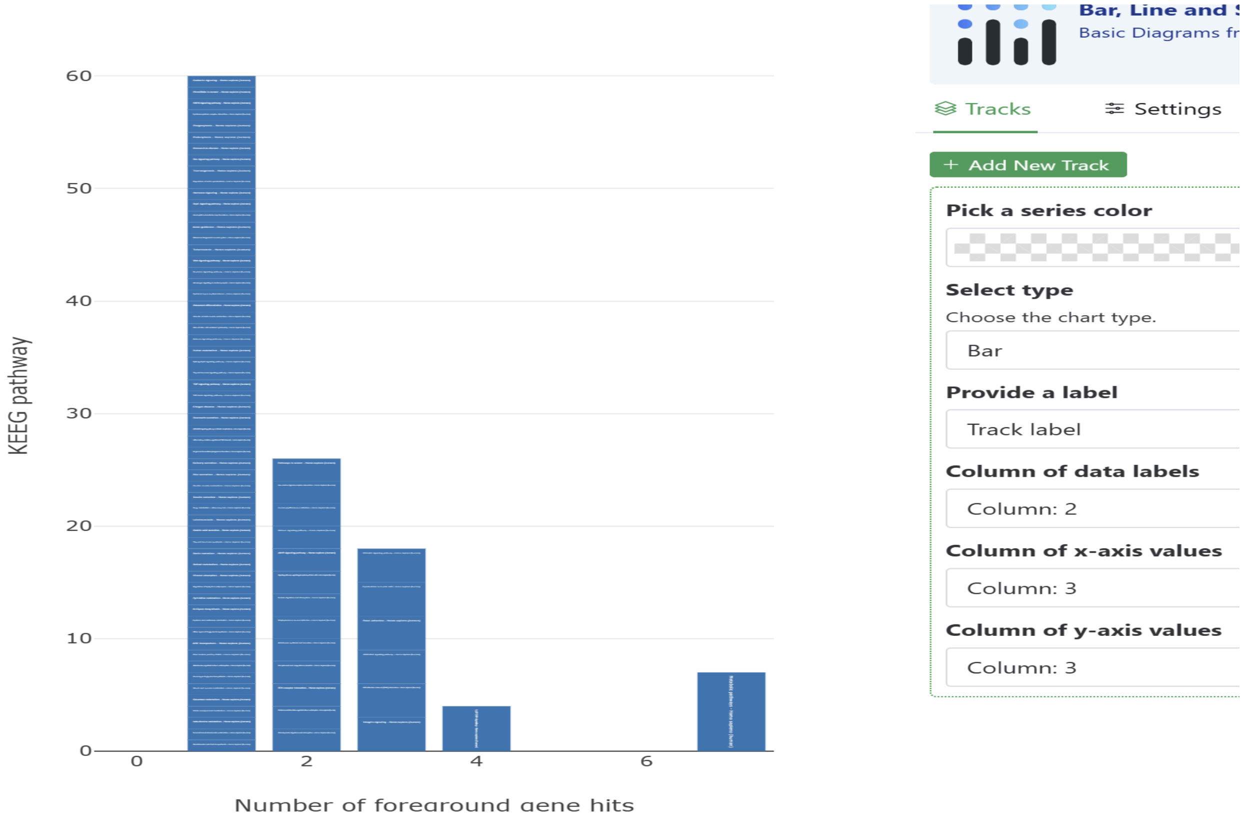

#### 192: KEGG ORA on dataset 191: pathway results

ok

373 lines 9 columns

format **tabular** database **hg38** size **30 KB**

Preview | Visualize | Details | Edit

| pathway_id | pathway_name | foreground_hits | pathway_genes_in_universe | foreground_genes_in_universe | universe_genes | |
|---|---|---|---|---|---|---|
| hsa04270 | Vascular smooth muscle contraction - Homo sapiens (human) | 2 | 135 | 15 | 9423 | 0. |
| hsa04080 | Neuroactive ligand-receptor interaction - Homo sapiens (human) | 3 | 371 | 15 | 9423 | 0.0 |
| hsa04360 | Axon guidance - Homo sapiens (human) | 2 | 184 | 15 | 9423 | 0.03 |
| hsa04614 | Renin-angiotensin system - Homo sapiens (human) | 1 | 23 | 15 | 9423 | 0.0 |
| hsa04950 | Maturity onset diabetes of the young - Homo sapiens (human) | 1 | 26 | 15 | 9423 | 0.0 |
| hsa00592 | alpha-Linolenic acid metabolism - Homo sapiens (human) | 1 | 27 | 15 | 9423 | 0.0 |
| hsa04081 | Hormone signaling - Homo sapiens (human) | 2 | 219 | 15 | 9423 | 0.0 |
| hsa00591 | Linoleic acid metabolism - Homo sapiens (human) | 1 | 31 | 15 | 9423 | 0.0 |
| hsa04014 | Ras signaling pathway - Homo sapiens (human) | 2 | 239 | 15 | 9423 | 0.0 |
| hsa05033 | Nicotine addiction - Homo sapiens (human) | 1 | 41 | 15 | 9423 | 0.06 |

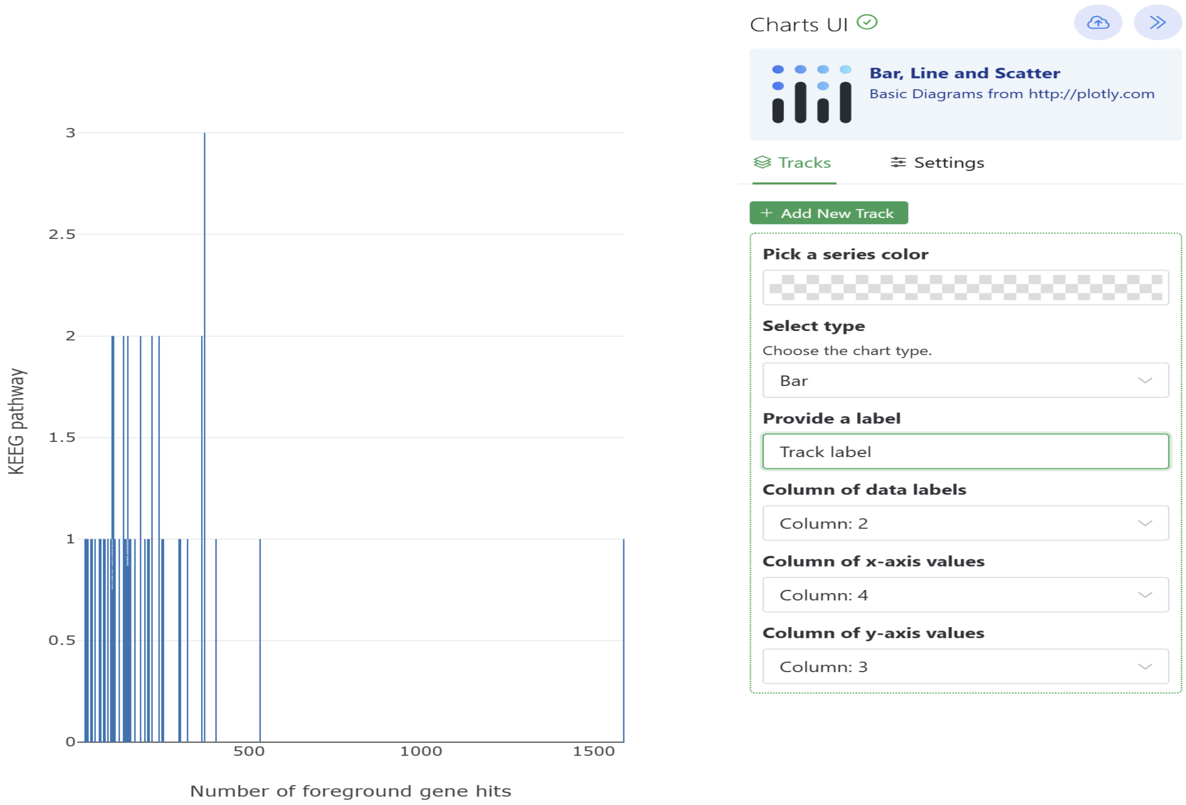


Figure 18,19 .KEGG Pathway Enrichment Results Both Upregulated & Downregulated.

| Gene set | KEGG result | Interpretation |
|---|---|---|
| Upregulated | 0 FDR-significant pathways | Nominal associations did not survive correction; minimum recorded adjusted p ≈ 0.71393 |
| Downregulated | 0 FDR-significant pathways | No pathway remained significant after correction; top adjusted p values were 1.0 |

Table 8. KEGG pathway enrichment summary.

No pathways were significant at an adjusted p-value cutoff of 0.05 in the original KEGG overrepresentation study. About 0.71393 was the lowest corrected p-value in my reported data. Consequently, I would still have no meaningful paths even if I changed the cutoff to 0.10 or 0.20. However, the input gene list and pathway results might modify if I redo the enrichment using a different, scientifically supported DEG selection threshold.

#### 8. Integrated biological interpretation

When all of these findings are combined, it can be said that the selected breast tumors and normal samples exhibit distinct transcriptional signatures. The idea that the examined samples have different transcriptomic profiles within the current data set is supported by the existence of 120 significant DEGs and the capacity to separate these groups using a heatmap.

It seems that the functional enrichment results are more selective. The GO Cellular Component *cell periphery* term was associated with the statistically verified enrichment in the case of upregulated DEGs. It indicates that cellular boundaries or cell-associated structures are somehow connected to some of the upregulated genes in the tumor group. Nevertheless, this analysis is unable to establish a causal connection between the differences found and tumor biology.

Since it relates to the boundaries of the current gene set, the lack of significant KEGG enrichment after correction is another finding that should be taken seriously. The number of DEGs, the consistency of changes, the mapping procedure, pathway coverage, or issues with multiple testing could all contribute to the lack of enrichment. This result must be presented rather than being changed to nominal p-value interpretation.

The original GSE210787 study should be taken into account when evaluating the biological context of the current dataset. Huang et al. (2024) reported the discovery of tumor-specific circular RNAs that produce cryptic antigenic peptides and their connection to anti-tumor immune responses. Since the current study focuses on traditional gene-level differential expression between the selected tumor and normal RNA-seq samples, it cannot be used to prove the mentioned immune mechanisms.

**9. Problems encountered and their resolution**

| **Issue** | **How it was handled** |
|---|---|
| Local STAR genome indexing stalled during junction insertion | The workflow was moved to Galaxy following the mentor's recommendation; STAR alignment completed successfully there. |
| 24 FASTQ files needed organization as paired-end samples | The datasets were assembled into a Galaxy paired-end collection containing 12 sample pairs. |
| Ensembl version suffixes caused identifier compatibility problems | Version suffixes were removed before enrichment and identifier conversion. |
| Some Ensembl IDs did not map to Entrez IDs | Unmapped entries were removed before KEGG ORA. |
| g:Profiler did not produce the expected KEGG terms | KEGG analysis was completed using a dedicated KEGG ORA workflow. |

Table 9. Problems encountered and their resolution.

**10. Final project outputs**

| Output | Galaxy / project reference | Purpose |
|---|---|---|
| Complete DESeq2 result | #145 | Full differential-expression statistics |
| Downregulated DEG list | #153 | 51 downregulated genes |
| Upregulated DEG list | #154 | 69 upregulated genes |
| Volcano plot | #155 | Effect size and significance visualization |
| Heatmap | #165 | Expression pattern of 120 significant DEGs |
| Downregulated GO enrichment | #180 | Functional enrichment results |
| Upregulated GO enrichment | #182 | Functional enrichment results |
| Upregulated KEGG results | #187 | KEGG ORA output |
| Downregulated KEGG results | #192 | KEGG ORA output |
| Final annotated DEG list | #200 | Consolidated DEG information |

Table 10. Final outputs retained for the project report.

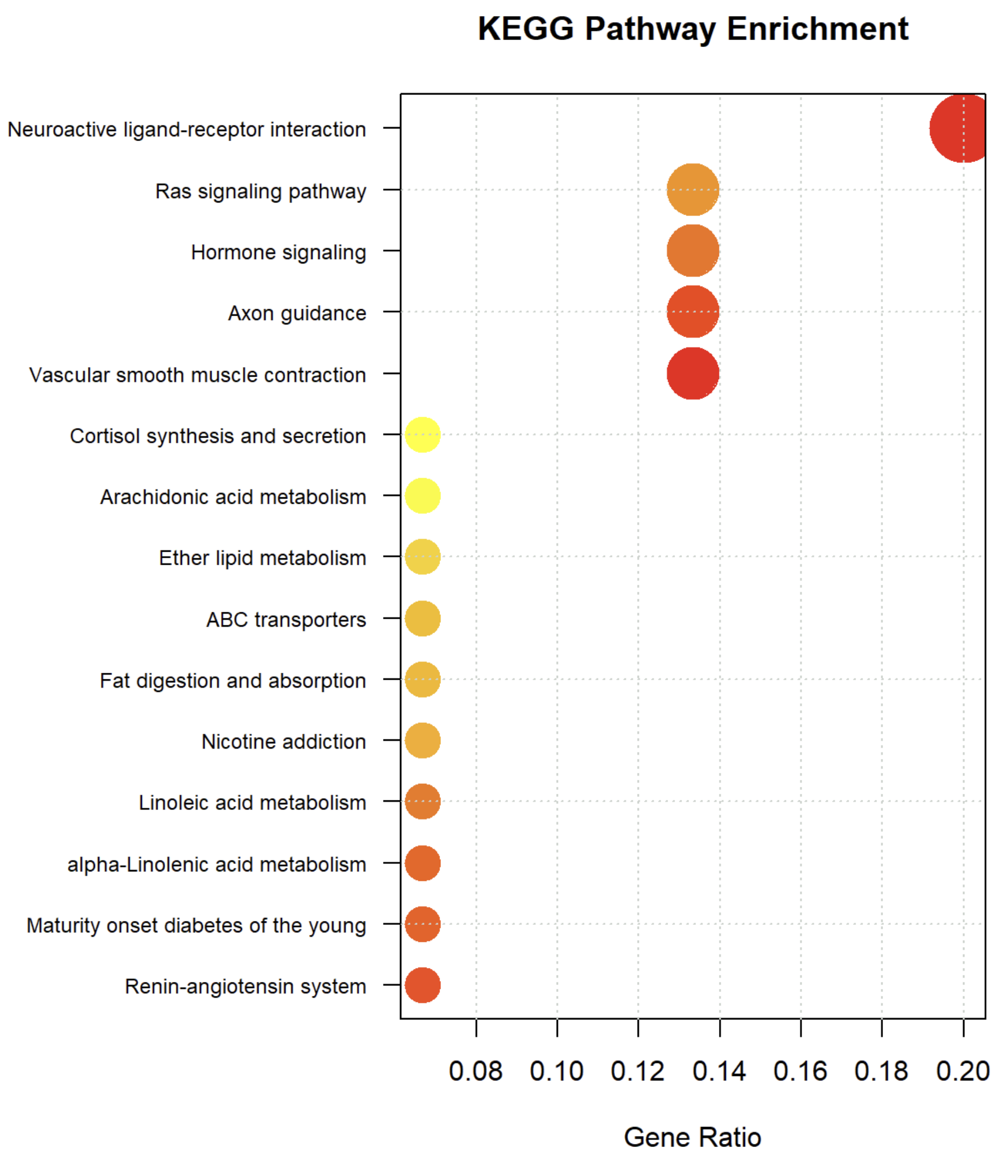


Figure 20. KEGG  Pathway Enrichment Results.

### CONCLUSION & FUTURE SCOPE

#### 1. Conclusion

Using the publicly accessible *GSE210787 dataset*, this project successfully finished an RNA-seq differential expression analysis comparing breast tumor and normal tissue. From data retrieval and quality assessment to read trimming, reference preparation, splice-aware alignment, gene-level quantification, differential expression analysis, visualization, and functional enrichment, the analysis followed a full workflow.

Using the predetermined criteria, *120 genes—69 upregulated and 51 downregulated*—were found to be differentially expressed. An overview of the correlation between the size of changes in gene expression and their statistical significance was given by the volcano plot. Additionally, the VST-based heatmap demonstrated distinct variations in the expression patterns of the normal and tumor samples.

Results from functional enrichment analysis were more constrained. Among the upregulated genes, GO enrichment revealed one statistically significant Cellular Component term, cell periphery; however, no significant GO terms were discovered for the downregulated genes. After adjusting for multiple testing, none of the nominal pathway associations revealed by KEGG analysis remained statistically significant. As a result, rather than being regarded as conclusive proof of biological enrichment, these pathway results were interpreted cautiously.

Learning how to stick to the overall analysis plan while making adjustments for computational constraints was one of the project's most useful practical components. Rather than altering the intended alignment method, the analysis was transferred to Galaxy when it became impractical to generate the STAR reference index locally due to computational requirements. This made it possible to preserve the initial methodological approach. while making the workflow more practical and manageable.

#### 2. Future scope

- When appropriate biological material is available, validate specific DEGs experimentally using an independent technique like quantitative RT PCR.
- To determine whether the detected expression changes are repeatable, repeat the differential-expression analysis in a separate breast cancer cohort.
- Because the source study has a significant circular-RNA component, look into changes at the transcript or isoform level, alternative splicing, and circRNA-specific signals.
- When appropriate, combine differential expression with other layers like proteomics, immune-cell infiltration, mutation data, or publicly accessible clinical data.
- When a larger or independently validated gene set becomes available, use ranked-gene approaches or other databases for pathway analysis.
- Instead of depending solely on enrichment-level summaries, investigate individual high-effect genes and potential regulatory mechanisms.
- Prior to making mechanistic or clinical claims, conduct independent validation and biological experiments.


**End of report.**
